# Small transcriptional differences among cell clones lead to distinct NF-κB dynamics

**DOI:** 10.1101/2021.12.07.471485

**Authors:** Cise Kizilirmak, Emanuele Monteleone, José Manuel García-Manteiga, Francesca Brambilla, Alessandra Agresti, Marco E. Bianchi, Samuel Zambrano

**Affiliations:** School of Medicine, Vita-Salute San Raffaele University, Milan, 20132, Italy; Division of Genetics and Cell Biology, IRCCS San Raffaele Scientific Institute, Milan, 20132, Italy; Center for Omics Sciences. IRCCS San Raffaele Scientific Institute, Milan, 20132, Italy

## Abstract

Transcription factor dynamics is fundamental to determine the activation of accurate transcriptional programs and yet is heterogeneous at single-cell level, even between very similar cells. We asked how such heterogeneity emerges for the nuclear factor κB (NF-κB), whose dynamics have been reported to cover a wide spectrum of behaviors, including persistent, oscillatory and weak activation. We found that clonal populations of immortalized fibroblasts derived from a single mouse embryo display robustly distinct dynamics upon tumor necrosis factor α (TNF-α) stimulation, which give rise to differences in the transcription of NF-κB targets. Notably, standard transcriptomic analyses indicate that the clones differ mostly in transcriptional programs related with development, but not in TNF-α signaling. However, by combining transcriptomics data and simulations we show how the expression levels of genes coding for proteins of the signaling cascade determine the differences in early NF-κB activation; differences in the expression of IκBα determine differences in its persistence and oscillatory behavior. The same analysis predicts inter-clonal differences in the NF-κB response to IL-1*β*. We propose that small (less than twofold) differences at transcript level can lead to distinct transcription factor dynamics in cells within homogeneous cell populations, and all the more so among different cell types.

## INTRODUCTION

Cells are able to provide precise transcriptionally-mediated responses to the complex mixture of external and internal stimuli to which they are subjected (Milo and Phillips, 2015). In this context, a key role is played by transcription factors (TFs), proteins that are “activated” upon stimuli and selectively trigger the expression of target genes coding for proteins required for an adequate response. The activation of several TFs is primarily mediated by their nuclear accumulation and is tightly regulated by other players within their genetic regulatory circuit, whose design contributes to providing a specific transcriptional output given a certain input (Alon, 2007). The nuclear accumulation of TFs is dynamic and can be oscillatory, as shown first for circadian rhythms in response to the day/night cycle (Patke et al., 2020) and the cell cycle (Ferrell et al., 2011); oscillations were then discovered for a wide variety of TFs (Levine et al., 2013). The emerging view is that such dynamics is not merely a by-product of the regulatory mechanisms of the TFs, but that it has a functional role in gene expression (Purvis and Lahav, 2013) and impacts a wide array of cellular process, e.g. determining cell fate (as for p53, (Purvis et al., 2012)), the response to mechanical cues (as for YAP/TAZ, (Franklin et al., 2020)) or the speed of the segmentation clock during embryo development (as for Hes7, (Matsuda et al., 2020)). Of note, single cell measures show consistently a high degree of heterogeneity in TF dynamics within a population, which yet is compatible with the TFs’ ability to provide an accurate transcriptional output given an input (Selimkhanov et al., 2014).

The NF-κB system is a paradigmatic example of the dynamic nature of TF activation (Aqdas and Sung, 2023; Kizilirmak et al., 2022). NF-κB is a family of dimeric TFs that plays a central role in innate and adaptive immune responses (Hayden and Ghosh, 2008; Natoli and Ostuni, 2019); dimers including the monomer p65 have the strongest transcription activating potential (Schmitz and Baeuerle, 1991) and are involved in the canonical pathway (we’ll refer to such dimers as NF-κB in what follows). NF-κB is kept in the cytosol bound by its inhibitors IκB, which are degraded upon external inflammatory stimuli such as the cytokine Tumor Necrosis Factor alpha (TNF-a) and are themselves NF-κB transcriptional targets (Hoffmann et al., 2002). It was immediately evident that this system of negative feedbacks could lead to oscillations in the nuclear concentration of NF-κB upon stimulation, as confirmed by an ever-growing list of live cell imaging studies (Nelson et al., 2004; Sung et al., 2009; Tay et al., 2010; Zambrano et al., 2014a). NF-κB nuclear localization dynamics (in short, NF-κB dynamics) have the potential to discriminate between ligand dose (Zhang et al., 2017) and type (Adelaja et al., 2021; Martin et al., 2020) and determines target gene expression (Ashall et al., 2009; Lee et al., 2014; Sung et al., 2009; Tay et al., 2010) in a functionally relevant way (Zambrano et al., 2016). However, terming such dynamics “oscillatory” is somewhat simplistic: the dynamics can be qualitatively quite different between cell types, ranging from sustained oscillations with a period of 1.5 hours for 3T3 cells (Kellogg and Tay, 2015), damped oscillations for mouse embryonic fibroblasts (Zambrano et al., 2016), persistent nuclear localization for RAW or 3T3 cells upon lipopolysaccharide (LPS) (Lee et al., 2009; Martin et al., 2020; Sung et al., 2014) and non-oscillatory for HeLa cells (Lee et al., 2014). Even within a population of cells of the same type, individual cells display dynamical heterogeneity and qualitatively different dynamics, with cells that oscillate and cells that do not (Nelson et al., 2004; Zambrano et al., 2014a). How different NF-κB dynamics arise, in particular within fairly homogeneous populations, is far from being clear and yet distinct NF-κB dynamics are crucial for cell specificity and can have important functional consequences, for example in the cell’s life-death decisions (Lee et al., 2016), its epigenetic state (Cheng et al., 2021) and even for the cell’s commitment to differentiation (Kull et al., 2022). Understanding how different dynamics emerge can shed light on how the NF-κB system uses dynamics in a cell-specific way to produce a desired output given a certain input, and more in general on the mechanisms by which cells within a population produce distinct TF dynamic responses to stimuli.

Here, we focused on immortalized GFP-NF-κB mouse fibroblasts (MEFs) derived from a single embryo. Yet, in these cells qualitatively and quantitatively different dynamics can be observed (De Lorenzi et al., 2009; Sung et al., 2009; Zambrano et al., 2014a; Zambrano et al., 2016). To determine the mechanisms responsible for such dynamical variations, we isolated several clones from the original cell population of MEFs and found that each clonal population has a distinct and heritable NF-κB dynamics upon TNF-α. We focused on three archetypical clones with persistent, oscillatory and weak responses, that show correlation with their transcription levels of NF-κB target genes. Cells within each clone are genetically and epigenetically identical. Instead, different clones share the same genome with sparse genetic variations accumulated during in vitro cultivation and immortalization, while their epigenetics instead shall reflect their heterogeneity in identity according to the anatomical site of origin (Lynch and Watt, 2018), as we confirmed through RNA-sequencing. We develop a framework for transcriptomic-constrained numerical estimation of the number of NF-κB activating complexes formed upon TNF-α or IL-1*β*, and we show that it is predictive of the relative strength of the early response to both inflammatory stimuli. Differences in expression of genes in the NF-κB signal transduction pathway, combined with small but robust transcriptional differences in the expression of the key negative feedback IκBα, explain the differences in the persistence of NF-κB activation in the clones. Indeed, we show how interfering with the expression of the repressor IκBα can make cells switch from a sharp response to a more persistent nuclear localization dynamics, as predicted by our mathematical model.

Taken together, our results show that small differences in the expression levels of the genes of the NF-κB signaling pathway can produce distinct responses in cells, even from quite homogeneous cell populations. This could contribute to explain how multicellular organisms produce selective and specialized NF-κB-mediated response to inflammatory stimuli. Furthermore, our framework can be used to investigate the origin of signaling variability across cells for other signaling pathways in other cell populations.

## RESULTS

### 1. Clonal populations of fibroblasts derived from a single embryo display distinct NF-κB dynamics upon TNF-a

Different NF-κB dynamics have been reported in different cell types, so here we decided to focus on a relatively homogeneous cell line: immortalized mouse embryonic fibroblasts (MEFs) derived from a single homozygous GFP-p65 knock-in embryo ((De Lorenzi et al., 2009) and **Methods**). We stimulated our GFP-p65 MEF population with 10 ng/ml TNF-α and quantified the nuclear to cytosolic intensity (NCI) of NF-κB using our established method of live cell imaging ((Zambrano et al., 2014a; Zambrano et al., 2016), and **Methods**). As previously reported (De Lorenzi et al., 2009; Sung et al., 2009; Samuel Zambrano et al., 2014a; Zambrano et al., 2016) the response of these MEFs upon TNF-α is heterogeneous (**Figure 1A** and **Movie S1**). Most cells display oscillatory peaks of nuclear localization while others display a non-oscillatory dynamics (**Figure 1B**), a kind of dynamics also referred to as “persistent activation” and similar to the one reported for macrophages (Cheng et al., 2021; Sung et al., 2014) and fibroblasts (Lee et al., 2009) upon LPS stimulation. We can indeed analyze NF-κB dynamics of hundreds of cells (see **Methods** and **Figure S1A**) and plot them to form a “dynamic heatmap” which captures the dynamics across the cell population (**Figure 1C**).

**Figure 1.**
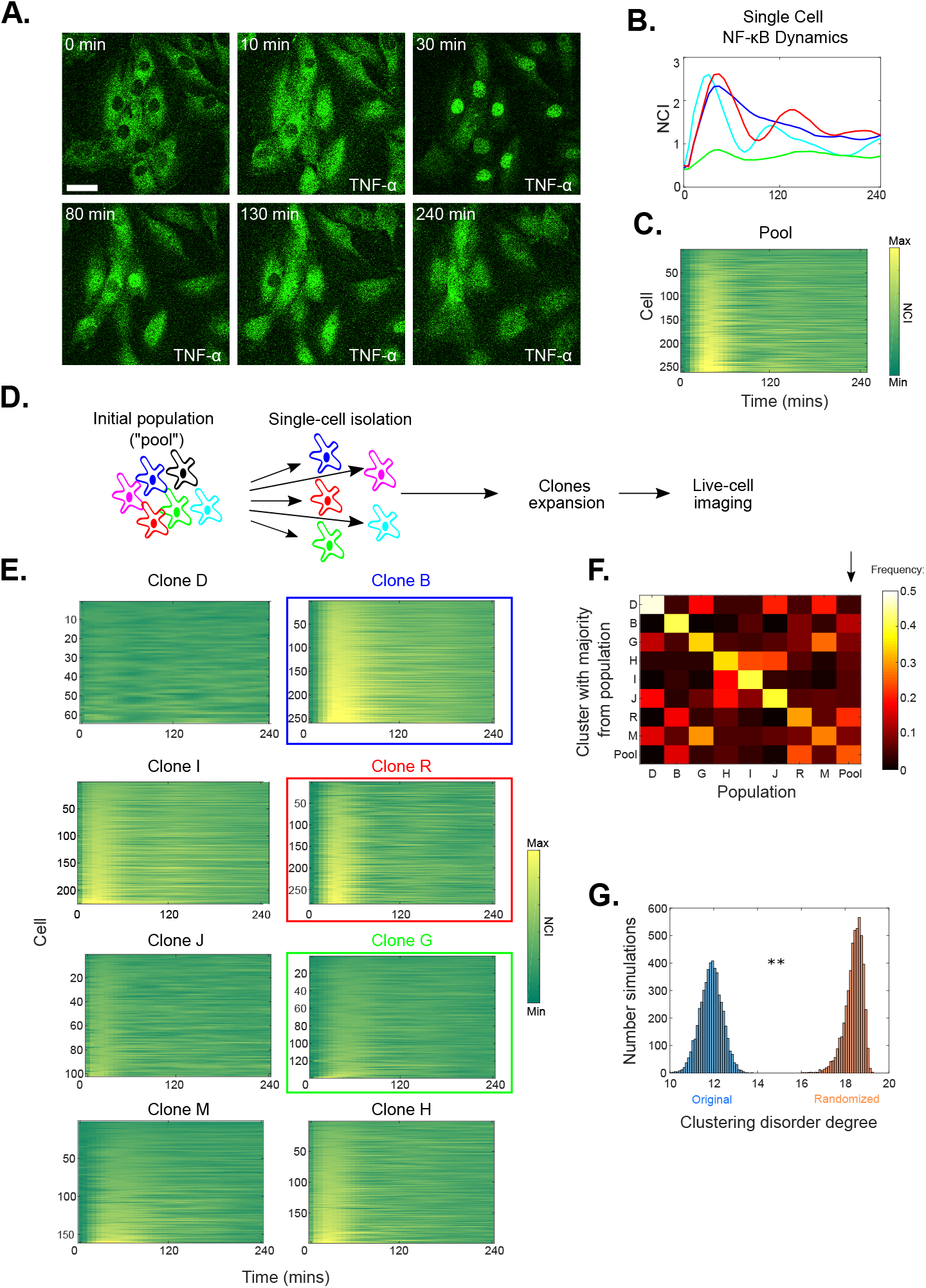
Clonal populations derived from a population of MEFs derived from a single embryo have distinct dynamics upon TNF-α. **A.** Representative images of the heterogeneous response to 10 ng/ml TNF-α of our initial population of MEFs, the pool. Scale bar indicates 20 μm. **B**. Heterogeneous NF-κB dynamic profiles of four cells from our MEFs population including weak (green), oscillatory (red) and persistent (blue) responses to 10 ng/ml of TNF-α. **C**. The dynamic heatmap of the pool, which represents the NF-κB dynamics of hundreds of cells sorted by their maximum NCI value. **D**. Single cell cloning strategy of our initial population of MEFs (referred to as “pool”). Clones were expanded and used for live cell imaging. **E**. Dynamic heatmaps of eight clones isolated from our population (Clones B, R and G are highlighted). **F**. The color-coded plot shows co-clustering probability of NF-κB dynamic profiles of each population considered (8 clones and the pool) based on an unsupervised k-means clustering. **G**. Distribution of the values of an entropy-like disorder parameter calculated for each realization of the stochastic clustering for the original dataset versus a randomized one. The distributions do not overlap in 500 realizations of the stochastic clustering for the original and the randomized datasets, p<2·10^−3^.

Starting from our original population of MEFs (that we refer to as the “pool” in what follows) we generated 17 clonal populations following standard procedures (**Figure 1D** and **Methods**). Eight clonal populations were imaged upon stimulation with 10 ng/ml TNF-α. Of note, these populations cannot be properly referred to as sub-clonal since the original population (the pool) does not derive from a single clone; instead, the pool is the result of a number of passages from a population of primary MEFs (see **Methods**), analogously to the procedure used to generate 3T3 cells (Amand et al., 2016; Gapuzan et al., 2005). We expect a small degree of genetic heterogeneity due to somatic mutations (Milholland et al., 2017) in the embryo and in the replications during the passages (we address this in further detail below). We found that NF-κB dynamics of each clonal population upon TNF-α was markedly different, even if a certain degree of intra-clonal heterogeneity was observed (**Figure 1E**). To have an unbiased confirmation of this observation, we utilized an unsupervised stochastic clustering approach (see **Methods**), which grouped NF-κB dynamic profiles of the clones and the pool according to their shapes into 9 different clusters (**Figure S1B**). For each realization of the clustering, we obtain a histogram with the number of profiles of a given population falling in a cluster where a given clonal population dominates (**Figure S1C**); to visualize them we can plot them together in a colormap (**Figure S1D**). Both single realizations of this procedure (**Figure S1D**) and the average of many (**Figure 1F**) indicate that NF-κB dynamic profiles of cells of the same population are more likely to cluster together than with profiles of cells of other populations (elements in the diagonal have higher values). Interestingly, cells from the pool are present in all the clusters where some of the clones are the majority (see arrow, **Figure 1F**), indicating that the dynamics of each clonal population mirror that of subpopulations of different sizes within the pool; in particular, clone R is the one with higher overlap with the pool. An entropy-based clustering disorder degree parameter (see **Methods**) based on these probabilities gives significantly lower values when the profiles are assigned to the proper clonal population (they are more “ordered” by the clustering) than when they are randomly mixed and then stochastically clustered (**Figure 1G**), further indicating that NF-κB profiles within a clonal population are similar.

In sum, we can isolate clones from a population of MEFs derived from a single embryo that have distinct dynamical behaviors which mirror those of individual cells in the original cell population.

### 2. NF-κB dynamics and oscillatory behavior are clone-specific and yet heterogeneous within clonal populations

We then chose 3 clones with archetypical dynamics reminiscent of those observed in the literature and in cells in our original MEF population (**Figure 1B**): a clone with more persistent nuclear localization of NF-κB (clone B, blue), one more oscillatory with a first well-defined sharp peak (clone R, red) (in other words, a nuclear localization that decreases fast (Zambrano et al., 2020)) and a clone characterized by a low activation of NF-κB (clone G, green) (**Figure 1E**).

Clonal populations R, G, and B showed clearly distinct NF-κB dynamics even by direct inspection of their timelapses (**Figure 2A** and **Movie S2-S4**). The average NF-κB activation profiles (**Figure 2B**) show that clone B gets more activated than clone R, and clone R than clone G: we can say they follow the order relation B>R>G. Such differences were strikingly robust across replicated experiments (examples of replicates shown in **Figure S2A**) and were conserved also for higher TNF-α doses (100 ng/ml) (**Figure S2B**). Importantly, these differences were also conserved after many cell divisions and culture passages, even if the average response to TNF-α was weaker for all clones after 8 weeks in culture (**Figure S2C**).

**Figure 2.**
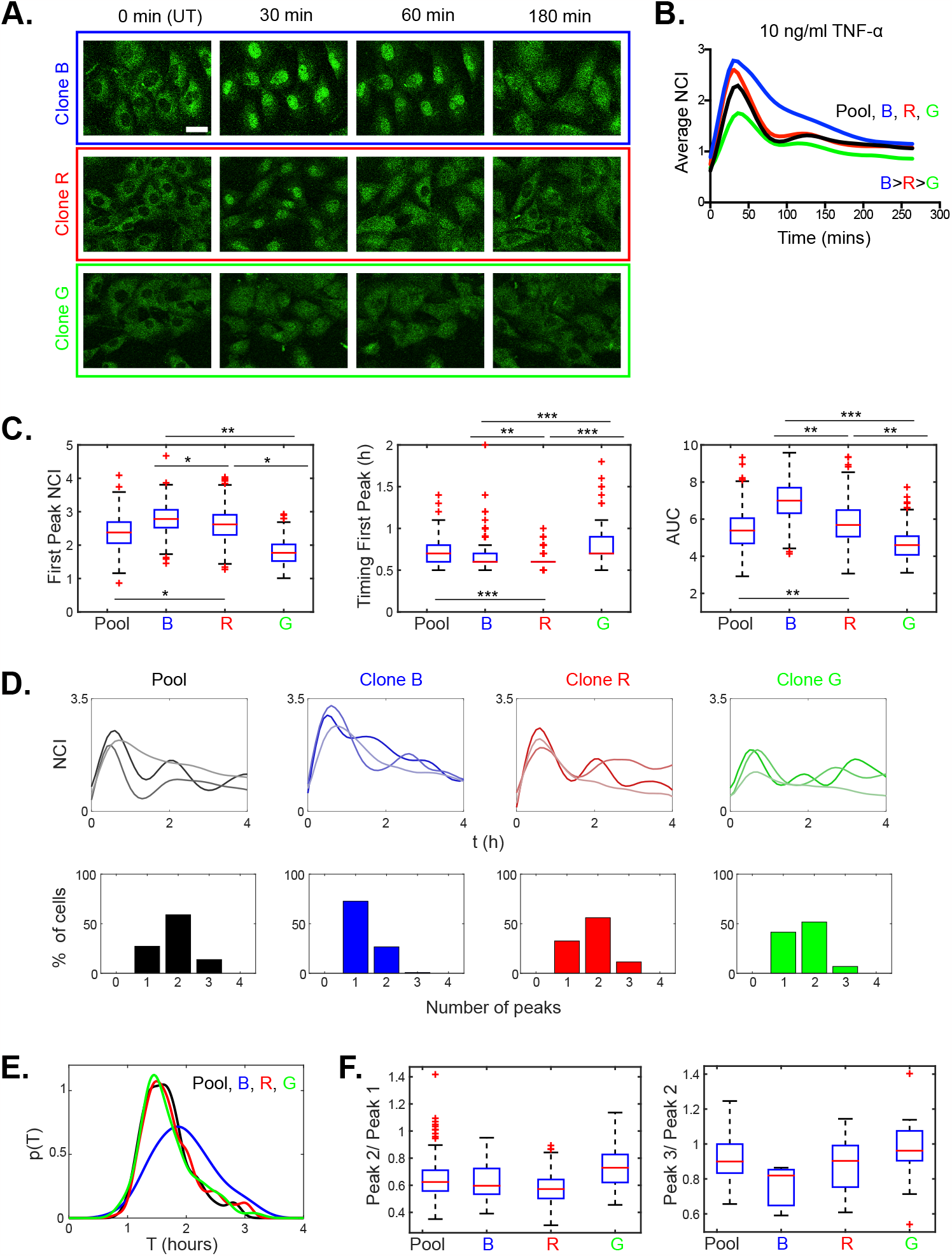
Clones B, R and G have distinct responses to TNF-α and are oscillatory to different degrees. **A.** Representative images from time-lapse movies for clones B, R and G upon 10 ng/ml TNF-α stimulation. Scale bar indicates 20 μm. **B**. NF-κB dynamic response of the clones and the pool to TNF-α as assessed by the average NCI. **C**. The boxplots show the dynamical features of the response to TNF-a: NCI value of the first peak, timing of the first peak and area under the curve (AUC) in each population. Pool and clones are always significantly different, significance is only displayed for comparisons between Pool and R. **D**. Examples of NCI profiles with 1, 2 and 3 peaks (top, lighter to darker colors) and frequency of the number of the oscillatory peaks observed (bottom) for each population in 4 hours. **E**. Periods of the oscillations computed as the inter-peak timing for each population. **F**. Ratios of the oscillatory peak values for each clonal population. In all panels *p<10^−2^, ** p<10^−3^, *** p<10^−4^, multiple comparisons through Kruskal-Wallis.

We then investigated the dynamics of each clone at single-cell level. The dynamic variability calculated through the coefficient of variation showed that clones’ dynamics have slightly less variability than the pool (**Figure S2D**). Next, we focused on typical single cell quantifiers of NF-κB dynamic response, such as the early NF-κB response of each cell (value of the first peak), its timing and the area under the curve (Tay et al., 2010; Zambrano et al., 2014a). We found that the values of such quantifiers are heterogeneous among cells of the same clone, although the differences are statistically significant between the clones and remarkably similar across replicates (**Figure 2C** and **Figure S2E**). The NF-κB response is typically higher for cells of clone B than for cells of clone R, which in turn have much higher peaks than cells of clone G (**Figure 2C**), so the B>R>G relation is preserved at single-cell level. Interestingly, clone G has the lower peak value in spite of having the higher basal levels of nuclear NF-κB (**Figure S2F**), resulting in an even lower fold change of nuclear NF-κB upon TNF-α with respect to the other clones (a key factor for gene expression (Lee et al., 2014)). The timing of the NF-κB response is equally prompt within our experimental resolution in clones B and R, but slightly slower in clone G (**Figure 2C**). The larger differences though are observed in the area under the curve (**Figure 2C**), which is much larger for clone B than for clone R, and is again the smallest for clone G. This captures well our observation that clone B has a more persistent NF-κB nuclear localization dynamics, while clone R displays a sharper response. All the quantifiers are significantly different between clones and the pool; in particular, they are different between pool and clone R, even if they do have the most similar TNF-α response (**Figure 2C**). We did also verify that all the differences above hold true when considering the absolute intensity of the nuclear signal (**Figure S2G**) instead of the nuclear to cytosolic intensity of NF-κB, and also when we manually segmented the cells’ nuclei (**Figure S2H**).

The differences between clonal populations could be related to factors such as cell cycle, which has been already shown to modulate the response to TNF-α (Ankers et al., 2016). Thus, we compared the cell cycle distributions of our populations (see **Methods, Figure S2I** and **Table 1**). Clone B and Clone G have very similar distributions, but they display the clearest differences in the dynamics of NF-κB response, suggesting that the cell cycle has a limited impact on their inter-clonal dynamic differences. Clones B and R are more different, so we applied a computational correction for the differences in the population fractions in each cell cycle phase (assuming that the stronger response in clone B is due to a higher fraction of cells in phase S than in clone R and correcting for this, details in **Methods**); even so, the differences were maintained (**Figure S2J**).

**Table 1:**
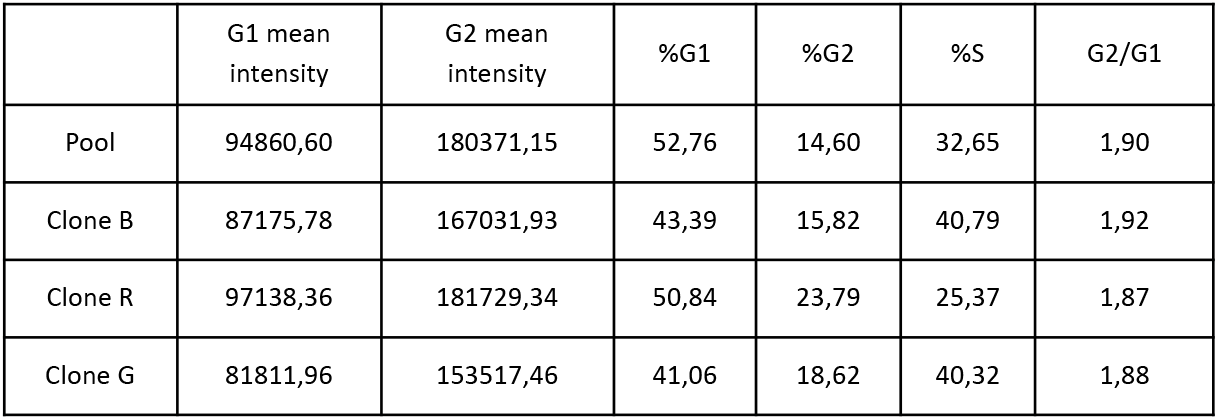
Percentages of cells in each cell cycle phase for each cell population.

Our single cell data does also provide us an interesting perspective on whether NF-κB signaling dynamics can be considered oscillating or not, a topic that has been subjected to discussion (Barken et al., 2005; Kellogg and Tay, 2015; Nelson et al., 2004; Zambrano et al., 2016). Oscillations are characterized by the presence of multiple peaks in the NF-κB dynamic profiles. We calculated the fraction of cells with 1, 2 or 3 oscillatory peaks within each clonal population upon treatment with 10 ng/ml TNF-α for 4 hours (**Figure 2D**) or for 12 hours (**Figure S2K**). We find that each clone contains a different fraction of cells that do have one peak in 4 hours (**Figure 2D**) but there are also cell fractions in each clonal population that have two or more peaks and can be considered “oscillatory”. Examples of each type of dynamics for each population are shown in **Figure 2D**; clone R could be considered the most oscillatory one. The period of the oscillations, computed as inter-peak timing, is heterogenous, close to 1.5 hours but slightly higher for clone B (**Figure 2E**). We also imaged our clones for 12 hours and computed the number of peaks, and only a minority of cells (but variable across clones) has 8 peaks or more and would be considered a “sustained oscillator” (**Figure S2K**). Clone R can hence be considered the more “oscillatory” one, but only in some cells we see sustained oscillations (see **Figure S2K**). Our data then show that being “oscillatory” is somehow a “fuzzy” phenotype and even clones derived from our MEF cell line can be considered oscillatory to different degrees. This compares with the qualitatively different dynamics observed for different cell types, that can go from cells that can oscillate in a sustained fashion for 20 hours and more (Kellogg and Tay, 2015) to our MEFs, where most of the cells oscillate in a damped way with fewer peaks (Zambrano et al., 2016), or HeLa cells that do barely oscillate (Lee et al., 2014). Consistently with the dynamic heterogeneity found in each cell population, we find variability in the peak values (**Figure 2F**) which is again maintained in experimental replicates (**Figure S2L**).

Overall, our data shows how clonal populations of MEFs derived from the same mouse embryo have distinct dynamical features at population level both in their early response to TNF-α and in the subsequent dynamics, which is oscillatory to a different degree within each population. Moreover, qualitatively and quantitatively heterogeneous dynamics can be found within very homogeneous populations, as our clones.

### 3. Clonal populations have distinct transcriptional programs and control of target gene expression upon TNF-a

To further investigate to what extent our clonal populations are different and to gain insights on how NF-κB controls target gene expression in each of them, we next performed RNA-sequencing. Our clones display quantitative differences in their average NF-κB dynamic response already after 1 hour TNF-α stimulation (**Figure 2B**). Therefore, we performed RNA-sequencing after 1 hour (**Figure 3A**). To gain statistical power, we generated 5 replicates per condition (see **Methods**). All replicates were of good quality with more than 10000 genes with CPM>1 (see **Methods** and **Figure S3A**).

**Figure 3.**
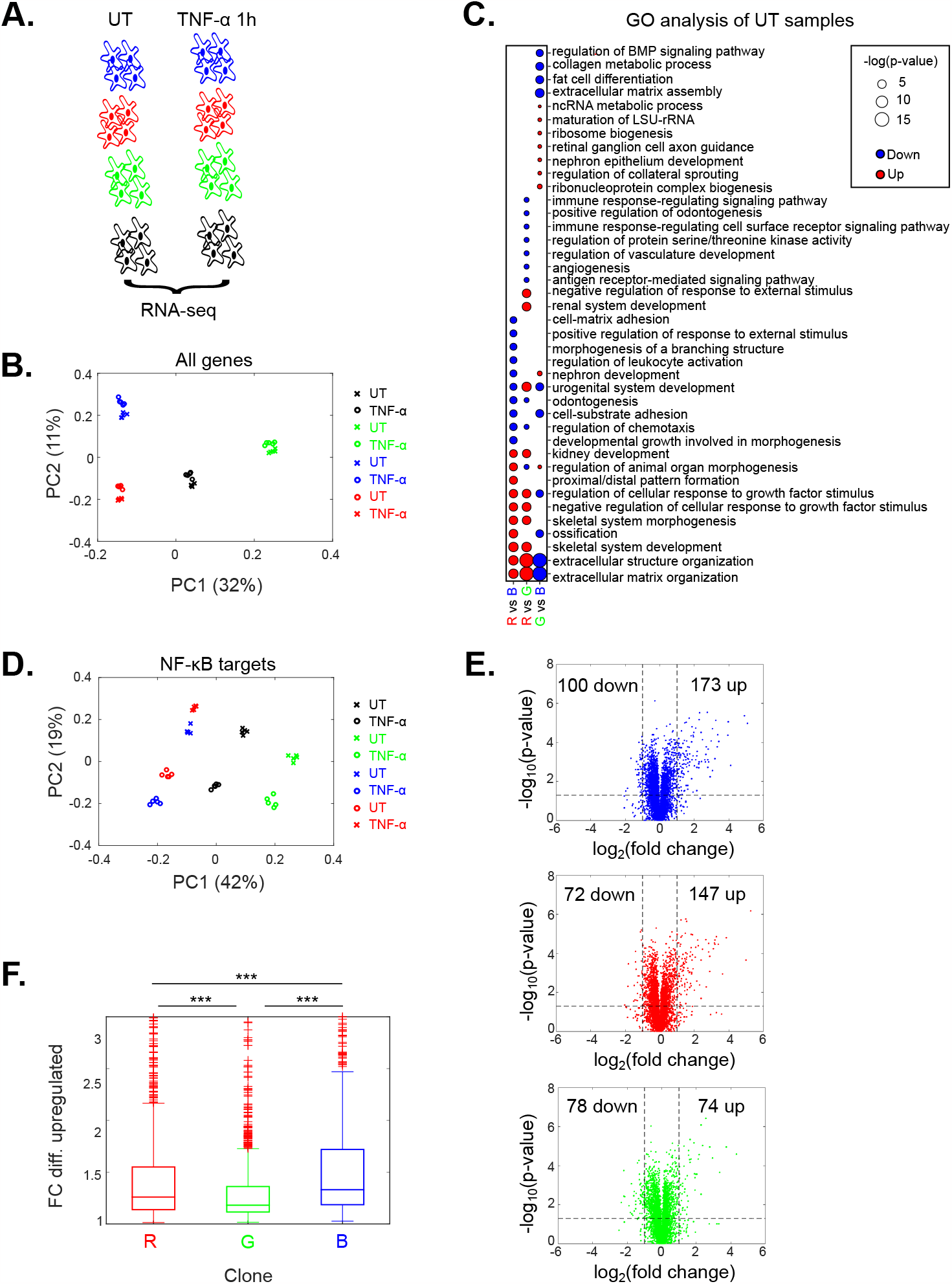
Clones B, R and G have distinct transcriptional programs. **A.** Scheme of the RNA-sequencing performed: 5 replicates per population, treated 1 hour with 10 ng/ml TNF-α or untreated (UT). **B**. PCA of the UT and TNF-α treated samples of our clones B, R, G and the pool, considering all genes (% of the variance indicated in parenthesis). **C**. Gene ontology (GO) analysis of the differentially expressed genes between the UT samples of the clonal populations. **D**. PCA considering only NF-κB targets (% of the variance indicated in parenthesis). **E**. Volcano plots of gene expression upon TNF-α for clone B, R and G (each dot is a single gene). **F**. Fold change expression of differentially upregulated genes in clones B, R and G. *p<10^−2^, ** p<10^−3^, *** p<10^−4^, multiple comparisons through Kruskal-Wallis.

RNA sequencing already provides insight on the high degree of genetic homogeneity of the populations, as expected from cells derived from the same embryo and subject to a low number of passages. The cells in the pool were immortalized following the 3T3 procedure from primary MEFs of the same embryo, a procedure that requires 20-30 passages (see **Methods**). Using an established reads mapping procedure (see **Methods**) we confirmed a frequency of single base substitution among clones (**Figure S3B**) compatible with the somatic mutation frequency found in cells of the same mouse (Milholland et al., 2017).

We then performed a PCA of our samples’ transcriptomes (see **Methods**) and found that samples from different clones do cluster in different groups (**Figure 3B**). Such neat clustering is also preserved in additional dimensions (**Figure S3C**). Of note, the apparent transcriptional divergence between our clones is small: when we performed our PCA including also public transcriptomic data from other tissues (see **Methods**), the samples from our populations cluster very closely together and far from the outgroup sample (**Figure S3D**); this might be partially due to technical and batch effects but the results indicate that the differences among our cell populations are small compared to the differences with and among samples of other tissues.

To get an unbiased insight on potential biological differences between our cell populations, we then looked at genes differentially expressed between clones that were untreated, which are in principle the most informative from the point of view of the cell’s identity. The categories enriched (see **Methods**) are mostly related to morphogenesis of different organs/tissues: epithelium, renal system and skeletal system, to cite a few (**Figure 3C**). This suggests that our clonal MEF populations conserve characteristics of the primary MEFs that were presumably already committed to different tissues or anatomical compartments (Lynch and Watt, 2018). The only categories reminiscent of NF-κB activation are “immune response-regulating signaling pathway”, “immune response-regulating cell surface receptor pathway”, “negative/positive regulation of response to external stimulus” and “regulation of leukocyte activation” (**Figure 3C**). Interestingly, a GO analysis performed for each clone against the pool shows that differences again are more related with developmental pathways; the fact that clone R and the pool have more similar responses to TNF-α than the remaining clones does not emerge from this analysis. (**Figure S3E**). To gain insights on the origin of the differences in dynamics across clones, we focused on the genes involved TNF-α signaling listed in the *wikipathways* database (see **Methods**) involved in TNF-α signaling. Of note, mutation calling did not highlight any differences in the mRNA sequences of these genes (see **Methods**). Although such genes cluster nicely by clones and treatments, overall they are more highly expressed in clone G (**Figure S3F**), so this classical unbiased analysis does not provide predictive insights on why the clones respond with different strength to TNF-α.

To gain insights on how NF-κB controls gene expression. First, we focused on 462 genes that we previously established are differentially expressed upon TNF-α stimulation (Zambrano et al., 2016). Interestingly, using this gene set the PCA now clusters samples both by treatment and by clone (see **Figure 3D** and **Methods**). This also applies when considering additional PCA dimensions (**Figure S3G**). We then had a closer look to the genes differentially expressed upon TNF-α for each clone (see **Figure 3E, Figure S3H** and **Methods**). Interestingly, the number of upregulated genes correlates well with the strength of the NF-κB response and satisfies the same order relation B>R>G (**Figure 3E**), and the fold change of the genes differentially upregulated also follow on average the B>R>G relation (**Figure 3F**). When performing gene set enrichment of upregulated genes, the recurrent categories include TNF-α signaling pathway and innate immune responses as expected (see **Methods** and **Figure S3I**), but with different degrees of overlap with the different categories, suggesting that the clones activate slightly different transcriptional programs upon TNF-α.

Our data show that the clones differ mostly for developmental identity and suggest that clonal differences in the strength of the NF-κB response are a key determinant of the expression level of NF-κB target genes upon TNF-α exposure. Clones differ transcriptionally mostly in GO terms not related to NF-κB, although genes belonging to the TNF-α signaling circuitry are also differentially expressed across the clones. However, this provides limited insight as to why the early NF-κB response follows the B>R>G order relation and why dynamics are different. Hence, we tried to gain additional mechanistic insight from our transcriptomic data by focusing on key players of NF-κB activation and describing mathematically the activation process.

### 4. Transcriptomic-constrained prediction of differences in the number of activating complexes between the clones explains the differences in their NF-κB responses to TNF-a

The activation of NF-κB upon TNF-α follows its interaction with its receptor, which leads to the formation of the so-called Complex I by the sequential association of different proteins (DeFelice et al., 2019; Hayden and Ghosh, 2008; Hsu et al., 1996; Lee et al., 2016; Wilson et al., 2009). These “activating complexes” formed upon TNF-α recruit and activate the IKK complexes (**Figure 4A**), which in turn determine the nuclear translocation of NF-κB (Cruz et al., 2021). We then reasoned that transcriptomic data might provide insights on whether different numbers of activating complexes are formed in the clones upon stimuli, resulting in the different NF-κB early responses observed (the first peak). We consider samples that are untreated, since activating complex formation is too fast to be impacted by transcriptional effects. We also assume that among our clones the ratios of mRNA levels mirror the ratios in protein abundances (see **Methods**). Although this could only be tested by a coupled proteomic and transcriptomic analysis of the clones, we find that this is the case for p65: mRNA abundance and fluorescence p65 intensity ratios are very similar among clones (**Figure S4A**).

**Figure 4.**
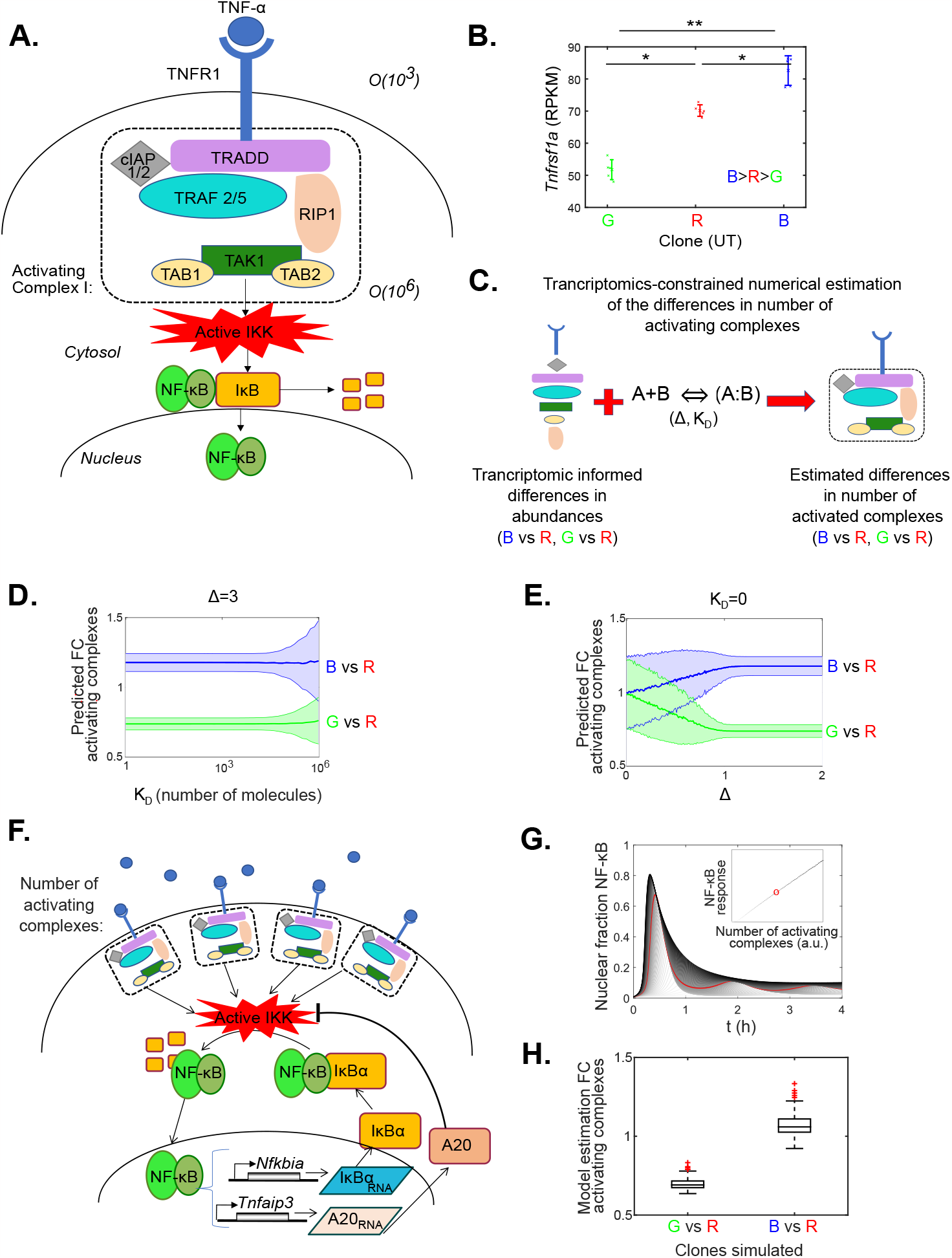
Transcriptomic-constrained predictions show the number of activating complexes formed upon TNF-α correlates with clones’ response. **A.** Scheme depicting the formation of the Complex I upon interaction of TNF-α with its receptor, leading to the IKK kinase complex activation. **B**. Expression level of the gene encoding for TNFR1 for the untreated clones, error bars display mean and standard deviations, each dot is a replicate. **C**. Scheme of the transcriptomic-constrained prediction scheme of the number of complexes formed for different values of Δ (differences in order of magnitude between copies of TNFR1 and rest of the proteins in the complex) and dissociation constant *K*_*D*_. **D**. Transcriptomic-constrained prediction of the fold change (FC) in the number of activating complexes formed upon TNF-α in clones B vs R (blue) and in G vs R (green) for the experimentally observed value Δ=3 and different maximum values of *K* _*D*_ considered in the simulations. Thick solid lines indicate the mean of the simulations, thin lines and shaded areas represent the standard deviation. **E**. Transcriptomic-constrained prediction of the FC in the number of activating complexes formed upon TNF-α in clones B vs R (blue) and G vs R (green) in the limit *K*_*D*_=0. Solid lines indicate the mean of the simulations, thin lines and shaded areas represent the standard deviation. **F**. Scheme of the NF-κB genetic circuit with main negative feedback regulators and including explicitly the number of activating complexes formed. **G**. Simulations of our mathematical model for a range of activating complex numbers and plot of the resulting NF-κB response for each value (inset), red line and red dot correspond to the same simulation. **H**. Result of estimating the FC in the number of activating complex needed in B vs R and G vs R to reproduce the differences in the NF-κB responses to TNF-α observed, for 250 different randomizations of the remaining model parameter values. *p<10^−2^, ** p<10^−3^, *** p<10^−4^, one-way Anova.

To investigate the possible cause of the differences in the response to TNF-α among clones, we elaborated the list “TNF-α to NF-κB” of genes coding for proteins forming the activating complex based on the existing literature (DeFelice et al., 2019; Hayden and Ghosh, 2008; Hsu et al., 1996; Lee et al., 2016; Wilson et al., 2009) (see **Figure 4A** and gene list in **Methods**) and found that the variation in their expression is quite small, less than 50% with respect to clone R (**Figure S4B**). Furthermore, single sample Gene Set Enrichment Analysis (Barbie et al., 2009) was performed for each of three clones to elucidate the enrichment of “TNF-α to NF-κB” list when compared to each other and to the original cell pool, but we did not find statistically significant differences (see **Methods** and **Figure S4C**). Taken together, standard bioinformatic analysis of the “TNF-α to NF-κB” list cannot explain the differences in the NF-κB responses observed between the clones.

We then moved to a more mechanistic description of the events leading to NF-κB activation. Activating complex formation can be considered as a sequential series of association (and dissociation) reactions of the proteins coded by genes of the “TNF-α to NF-κB”. Importantly, these proteins are upstream of NF-κB; proteins downstream might have an influence but for the sake of simplicity we do not consider their role in early responses. Of note, the total abundance of the activating complex will be limited by the less abundant component (it is not possible to have more copies of the complex (*A*: *B*) than the minimum of *A* and *B* abundances, see **Methods** and **Figure S4D**). In our “TNF-α to NF-κB” list, the less abundant protein is known to be the receptor TNFR1 (encoded by *Tnfrsf1a*) of which there are about 10^3^ copies, while the abundance of the remaining proteins in the list is about 10^6^ (Cruz et al., 2021; Hwang et al., 2015). Hence TNFR1 is the limiting factor in the formation of activating complexes and their number will be largely determined by its abundance (see **Methods**), irrespective of the relative abundances of other proteins. Interestingly, transcriptomic data shows that the expression of TNFR1 in untreated samples follows the order relation B>R>G (**Figure 4B**) which in should lead to a number of activating complexes following B>R>G, which correlates with the experimentally observed ranking in the early NF-κB response upon TNF-α.

To further test whether the order relation B>R>G would also hold when all the processes of association-dissociation involved in complex formation and all the differences in abundance of the proteins considered are taken into account, we devised a transcriptomic-constrained numerical approach to estimate the fold change (FC) differences in number of complexes between the clones (**Figure 4C**). We simulate the association reactions between the elements of the “TNF-α to NF-κB” list leading to the formation of the activating complex, each characterized by their dissociation constant *K*_*D*_, which we consider equal for each reaction among all the clones (see **Methods**). We assume again that ratios in protein abundances between clones match that of transcript abundances, as for p65 (**Figure S4A**). The average protein abundance of each element in the list across clones is chosen randomly in each numerical simulation, but taking into account the difference in orders of magnitude Δ between the abundance of the receptor (O(10^3^)) and the remaining proteins (O(10^6^)), that is Δ=3 (Cruz et al., 2021; Hwang et al., 2015). Numerical estimates (**Figure 4D**) predict an order relation B>R>G for a wide range of order of magnitudes of the *K*_*D*_ values, although variability increases for high *K*_*D*_, a situation where the number of complexes is low (see **Figure S4D** and **Methods**). We also evaluated the dependence of the result in Δ by performing simulations in the limit situation *K*_*D*_ =0, where the association between the proteins that form the complex is perfect, and for Δ values much lower than the experimentally observed ones: we find that the transcriptomic-constrained predictions of the number of complexes converge quickly to a B>R>G ranking (**Figure 4E**). Simulations in the *K*_*D*_-Δ parameter space confirm these trends (**Figure S4E** and **S4F**) and show that the B>R>G order relation of the number of activating complexes holds for a wide parameter range.

To test whether the transcriptomic-constrained predicted small differences in the number of activating complexes can explain the different observed responses we developed a simple mathematical model of NF-κB signaling based on previous ones (Zambrano et al., 2014b; Zambrano et al., 2016) (see **Methods** and **Figure 4F**). In our model, the number of activating complexes determines the activation of the IKK kinases, so the NF-κB response and the number of complexes correlate (Cruz et al., 2021) (**Figure 4G**). We can then use this model and estimate the FC in the number of complexes of B vs R and G vs R leading to changes in the early response. In doing so we take also into account the variations in NF-κB (p65) amount across clones (**Figure S4A**), which alone has a small impact (**Figure S4G**). An example of how that FC is computed is shown for a given parameter set (provided in **Table 3**) in **Figure S4H**. To assess in a more general way the FCs in the number of activating complexes that would be needed to recapitulate differences in the experimentally observed responses, we repeated this estimation for different randomizations of the initial parameters of the model (see **Methods** and examples in **Figure S4I**). The results (**Figure 4H**) indeed show that for most parameter sets small changes in the number of activating complexes are enough to reproduce the observed differences in the early response, with values within the range of those obtained from our transcriptome-constrained estimations (compare **Figure 4D, 4E** with **Figure 4H**).

Taken together, our numerical approach suggests that small differences in the transcription levels of the TNF-α receptor are responsible for the differences in the early NF-κB responses observed between the clones. Transcriptomic-constrained estimations and mathematical modeling show that the B>R>G relation in the number of activating complexes would hold for a large range of values of the parameters involved, and that differences in TNFR1 are enough to explain the differences in the NF-κB response between the clones.

### 5. Transcriptomic-constrained simulations predict differences in early NF-κB activation between the clones upon IL-1*β*

Since our transcriptome-constrained prediction of the number of activating complexes explain the differences in the early response between the clones to TNF-a, we asked if they could also explain the differences in the response to a different inflammatory stimulus. We focused on IL-1*β*, a cytokine that is known to activate NF-κB through a pathway only partially overlapping with that of TNF-a, and characterized by its own regulatory mechanisms and dynamical features (DeFelice et al., 2019; Martin and Wesche, 2002). We elaborated a list of “IL-1*β* to NF-κB” genes coding for proteins that are involved in the formation of the activating complexes arising upon interaction of IL-1*β* with its receptor (Cruz et al., 2021; Martin and Wesche, 2002) (elements shown in **Figure 5A**, gene list in **Methods**). The list now includes genes coding for the subunits of the dimeric receptor, *Ilr1r1* (coding for IL-1R1) and *Il1rap* (coding for IL-1R3). In this list, differences in expression across clones are small and typically do not exceed 2-fold (**Figure S5A**); ssGSEA finds a difference between clone B and the other clones, but not between the remaining ones (**Figure S5B**). For this signaling system it has been shown that the limiting factor are the dimeric receptor proteins IL-1R1 and IL-1R3, of which there are O(10^2^-10^3^) copies, Δ≳3 orders of magnitude less abundant than the remaining downstream proteins (Cruz et al., 2021; Hwang et al., 2015). Interestingly, we found that expression of *Il1rap* is very low compared to *Il1r1* (**Figure S5C**) and in the untreated samples follows the order relation B≳G>R (the differences between B and G are not significant, but in both it is more highly expressed than for R (**Figure 5B**)). Thus, the expectation is that the number of activating complexes formed by the clones should follow the order B≳G>R.

**Figure 5.**
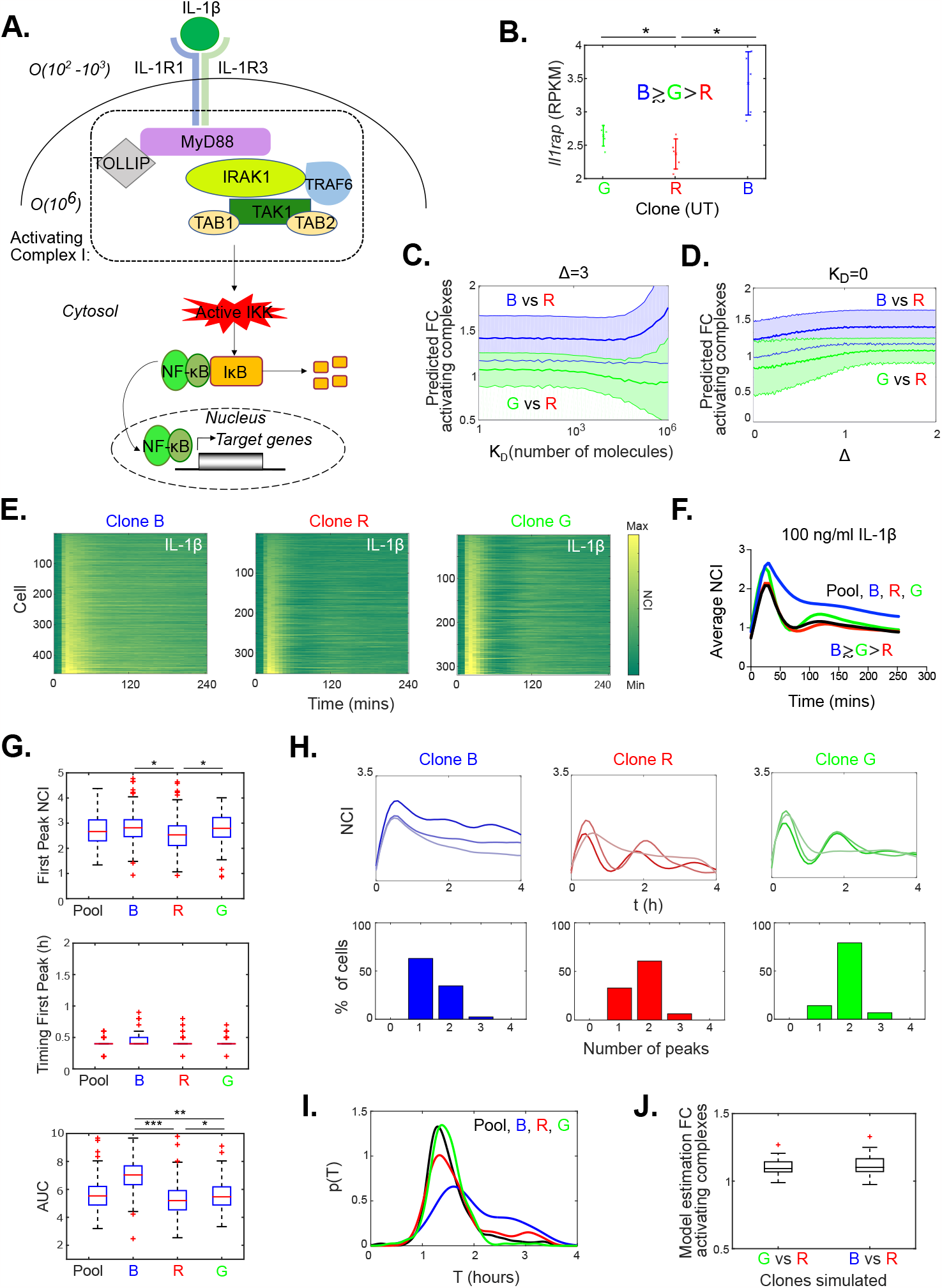
Transcriptomic-constrained simulations predict differences in the clones’ response to IL-1*β*. **A**. Scheme depicting the formation of the activating complex arising upon interaction of IL-1*β* with its dimeric receptor. **B**. Expression level of the gene encoding for IL-1R3 for the untreated clones, error bars display mean and standard deviations, each dot is a replicate. **C**. Transcriptomic-constrained prediction of the FC in the number of activating complexes formed upon IL-1*β* by clones B vs R (blue) and G vs R (green) for the experimentally observed value Δ=3 and different values of *K* _*D*_ considered in the simulations (the most abundant proteins can have up to 10^6^ copies in the simulations). Thick solid lines indicate the mean of the simulations, thin lines and shaded areas represent the standard deviation. **D**. Transcriptomic-constrained prediction of the number of activating complexes formed upon IL-1*β* by clones B vs R (blue) and G vs R (green) in the limit *K* _*D*_ =0. Solid lines indicate the mean of the simulations, shaded areas within thin lines represent the standard deviation. **E**. Dynamic heatmap of the responses of clone B, R and G to 100 ng/ml IL-1*β*. **F**. Average NF-κB response for the three clones and the pool upon IL-1*β* stimulation. **G**. The boxplots show the dynamical features of the response to IL-1*β*: value of the first peak, timing and area under the curve (AUC). **H**. Examples of NCI profiles with 1, 2 and 3 peaks (top, lighter to darker colors) and frequency of the number of the oscillatory peaks observed for each clonal population (bottom) in 4 hours upon IL-1*β*. **I**. Periods of the oscillations computed as the inter-peak timing for each population. **J**. Model estimation of the FC in the number of activating complex needed to reproduce the differences in the NF-κB responses to IL-1*β* experimentally observed in B vs R and G vs R, for different randomizations of the remaining parameter values. *p<10^−2^, ** p<10^−3^, *** p<10^−4^, multiple comparisons through Kruskal-Wallis.

We then performed our transcriptomic-constrained prediction of the FC variation in the number of activating complexes between clones for a wide variety of conditions (see **Methods**). Given Δ=3 we find that the relation B>G>R holds for a wide range of values of *K*_*D*_ (**Figure 5C**) but with a higher variability than for TNF-a, in particular for high *K*_*D*_, so B≳G>R would be more accurate. We also find that the estimations quickly converge to B≳G>R for Δ values smaller than the one experimentally verified (Cruz et al., 2021; Hwang et al., 2015) (**Figure 5D**). These trends are also confirmed when we analyze the *K*_*D*_-Δ parameter space (**Figure S5D** and **S5E**). Overall, this analysis suggests that the number of activating complexes upon IL-1*β* shall follow the relation B≳G>R, with B and G having similar NF-κB responses and both with a response stronger than R.

To test this prediction, we performed live cell imaging of the clones upon 100 ng/ml IL-1*β*. All clones respond, as described by others (Adamson et al., 2016; DeFelice et al., 2019), but again in a clone-specific way: in particular now clone G has a stronger response to IL-1*β* than to TNF-a, with an early NF-κB response (first peak value) similar to that of clone B (**Movies S5 and S6** and **Figure 5E**). Importantly, the average NF-κB response reflects the predicted order relation B≳G>R with clone B and G having similar first peak value (**Figure 5F**). We found these differences are also clear when considering single-cell data (**Figure 5G, S5F**), where clone G cells display overall a stronger response than clone R cells. However, clone B is the one with a higher response and a much higher AUC. Finally, upon IL-1*β* we find that the cells’ oscillatory phenotype is clone-specific and yet heterogeneous for each clone (**Figure 5H**), but different relative to TNF-a: more cells of clone G have at least two peaks (**Figure 5H**) and an oscillatory period T=1.5h, similar to clone R (**Figure 5I**). Oscillations are similar to those observed upon TNF-a, as described for other cell types (Adamson et al., 2016; DeFelice et al., 2019). Clone B remains the one with a less oscillatory phenotype (**Figure 5H**), suggesting that this behavior might be related with downstream regulators of NF-κB activity, an idea that we explore next. As with TNF-a, the parameters describing the oscillations are also heterogeneous across the clones (**Figure S5G**).

Finally, following the procedure described for TNF-α (see also **Methods**), we assessed to what extent the FC variation in the number of activating complex complexes predicted by transcriptomics can reproduce the observed differences in the early response (first peak). We used our mathematical model of NF-κB dynamics to estimate the FC required for different parameter sets (example for one parameter set shown in **Figure S4H**). Simulations show that for most parameters considered a variation in the number of activating complexes within the range predicted by our transcriptomic-constrained estimation (**Figure 5J**) reproduces the observed experimental differences in the early NF-κB responses between our clonal populations.

Taken together, our transcriptome-constrained modeling approach predicts that a small difference in the expression of the limiting factor for formation of the activating complex upon IL-1*β* (the IL-1R3 receptor subunit) would lead to the B≳G>R ranking, a prediction that is confirmed by live cell imaging experiments.

### 6. Differences in the expression levels of the main negative feedback underpin distinct NF-κB dynamics of clonal populations

Beyond differences in the first peak of the response, our clonal populations differ also at later time points. For the sake of simplicity we focus on the case of clone B having a more persistent NF-κB response -quantified by a higher AUC-as compared to clone R both for TNF-α (**Figure 2C**) and IL-1*β* (**Figure 5G**). However, our model estimates that by changing the NF-κB amount and the number of activating complexes to match the differences in the early responses for a wide variety of combinations of the remaining parameters only minor changes in the AUC are obtained, well below the experimental differences observed between clone B and R (**Figure 6A**, example of one calculation shown in **Figure S6A**).

**Figure 6.**
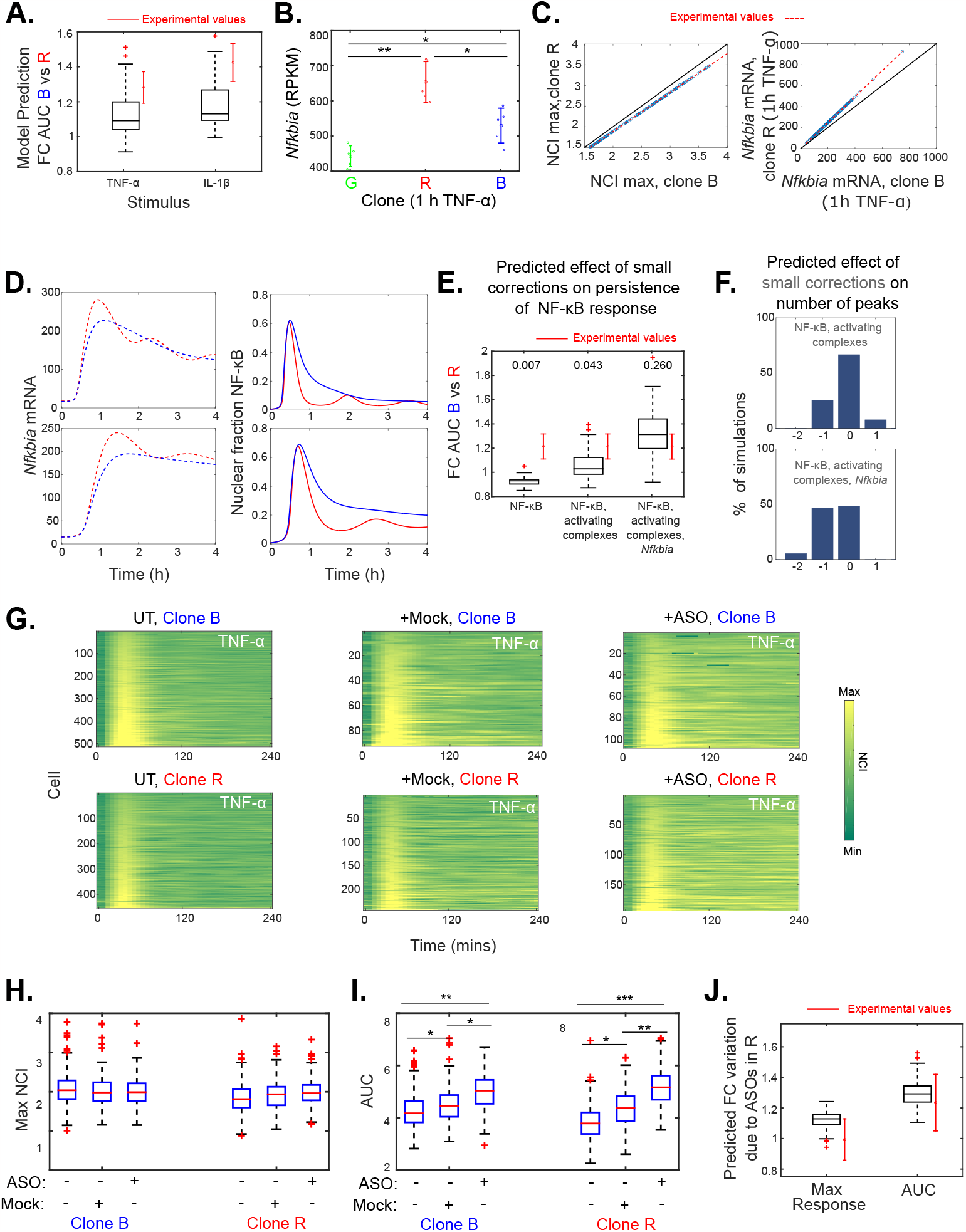
Differences in expression levels of NF-κB negative feedbacks can reproduce the observed dynamical differences between clones. A. Resulting change in the AUC of the clone B against clone R when fitting the number of activating complex needed to match the differences in their early NF-κB responses to TNF-α and IL-1*β* for different combinations of the remaining parameter values, and mean and standard deviation of experimental replicates (red). **B**. Expression level of the gene encoding for IκBα (*Nfkbia*) for clones after 1 hour of TNF-a, error bars display mean and standard deviations, each dot is a replicate. **C**. Maximum NCI response (left) and *Nfkbia* levels (right) of our transcriptionally constrained numerical simulations of clone R and clone B, each dot is a single simulation where parameters of clone R and clone B have been adjust so the ratios of these quantities match the experimental observed ones (dashed red). Solid black line is the identity. **D**. Exemplary pair of simulations with the imposed changes in *Nfkbia* levels shown in C (dashed) leading to a change in the dynamics (solid line) in the simulations of R versus B. **E**. Changes of the AUC resulting of correcting the following model parameters, following the experimental data: the amount of NF-κB, both the amount of NF-κB and the number of activating complexes to fit the NF-κB response, and the two previous combined with the amount of *Nfkbia*, for different randomization of the model parameters. Red errorbar represents the mean and standard deviation of changes in AUC observed in three experimental replicates. Values reported for p-value of Mann-Whitney test. **F**. Change in the number of peaks for simulations giving oscillatory profiles (at least 2 peaks), when only the number of activating complexes and the NF-κB amount are changed (top), and also when the amount of *Nfkbia* is changed (bottom). **G**. Representative dynamic heatmaps of clone B and R upon TNF-α for untreated, scrambled ASO (MOCK) and *Nfkbia*-targeting ASO treated cells. **H**. Quantification of the maximum response for untreated, MOCK treated and *Nfkbia* targeting ASO treated cells upon TNF-α. **I**. Quantification of the AUC for untreated, MOCK treated and and *Nfkbia*-targeting ASO treated cells upon TNF-α. **J**. Simulated effect of modulating *Nfkbia* as observed experimentally in the response and the AUC of different randomizations of the model’s parameters. Red errorbar represents the mean and standard deviation of changes in the AUC observed in experimental replicates. *p<10^−2^, ** p<10^−3^, *** p<10^−4^, multiple comparisons through one way Anova (panel E) and Kruskal-Wallis (remaining panels).

To explain differences in AUCs, we then decided to focus on the transcriptional differences in the negative feedbacks. We focused on IκBa, encoded by *Nfkbia*, the key gene in determining the shift between oscillating and persistent dynamics (Cheng et al., 2021; Hoffmann et al., 2002). With IκBa, we are considering relatively short-term dynamics, while the remaining negative feedbacks of the system (Ashall et al., 2009; Hoffmann et al., 2002) have a stronger effect on longer term responses and population heterogeneity (Paszek et al., 2010; Son et al., 2021). Moreover, the remaining negative feedbacks are expressed at much lower levels for UT and TNF-α treated samples (**Figure S6B**), with similar fold change for all clones (**Figure S6B**).

Our transcriptome data show that the expression of IκBα upon 1 hour TNF-α (when it peaks) follows a R>B>G pattern (**Figure 6B**) confirmed by RT-qPCR (**Figure S6B**), which correlates with the observed differences in the dynamics. To dissect in a general way if this difference, together with the previously described, is enough to recapitulate the experimentally observed differences in the dynamics among the clones, we performed experimentally constrained numerical simulations using our mathematical model. We used different randomized parameters for clone R, where we sequentially adjust the amount of NF-κB, together with the FC in the number of activating complexes to match the FC in the experimentally observed responses between clone B and R (**Figure 6C**, left), and the transcription rate of *Nfkbia* to fit the experimentally observed FC differences in the expression level of B respect to R (**Figure 6C**, right). This leads to different prototypical dynamics of B and R (**Figure 6D**), just by changing 3 parameters (see **Methods**). In **Figure 6E** we show the change on the AUC predicted by our model by applying corrections in the parameters that mimic the experimental differences between clone B and R, to hundreds of parameter sets. Of note, only the combination of three changes (in NF-κB amount, activating complex number and *Nfkbia* expression) results in a change in the AUC compatible with the difference observed between clone B and clone R (**Figure 6E**). We do not exclude that fine tuning also other parameters, including more feedbacks, would lead to a change closer to experimentally observed values. Interestingly, when we analyze the effect of each correction on the oscillatory profiles, we find that the modulation of *Nfkbia* is determinant in producing less oscillatory profiles (**Figure 6F**), something that can also be observed in the examples displayed in **Figure 6D**. This indicates that the experimentally constrained small differences in these model parameters, combined, are sufficient to explain experimental differences in the persistent and sharp responses of clone B and R, respectively, and why clone R has more oscillatory dynamics than clone B.

To experimentally test the effect of modulating IκBα levels in the dynamic response to TNF-α we took advantage of antisense oligonucleotide (ASO) technology (see **Methods**). The single-stranded synthetic 2’-deoxy-2’-fluoro-β-d-arabino nucleic acid (FANA) oligos were designed to be self-delivered to the cells and to be fully complementary to IκBα mRNA, inducing mRNA cleavage by RNAseH and reducing the synthesis of the IκBα protein (see **Methods**). In our pool, scrambled (MOCK) ASOs induced no change in the activation but a small change in the AUC, compared to the effect of *Nfkbia*-targeting ASOs (**Figure S6D**). Hence we repeated the approach with our clones B and R. When treated with *Nfkbia*-targeting ASOs, we indeed observed that the response of both clone B and clone R (**Movies S7** and **S8** respectively) was characterized by more persistent NF-κB nuclear localization; this indicates that a moderate but significant decrease on the mRNA levels of IκBα in both clones (**Figure S6E**) is enough to produce a qualitative change in the clonal dynamics (**Figure 6G**). Quantification of the dynamics shows that the maximum value of the NF-κB response is almost unaltered in ASO-treated cells in clone R (**Figure 6H**). However, when treated with *Nfkbia*-targeting ASOs, the area under the curve upon TNF-α increases for both clones (**Figure 6I**), and in particular for clone R, so that its previously described oscillating dynamics are modulated to be similar to that of clone B. This trend is reproducible in replicates (**Figures S6F, S6G**). The effect of the ASOs in clone R can be reproduced in numerical simulations for different randomization of the model parameters, where we only adjust *Nfkbia* expression levels following the experimentally observed changes (from **Figure S6E**) in expression (see **Figure 6J** and **Methods**). Of note, as for the pool (**Figure S6D**), scrambled ASOs (MOCK) have a smaller (but significant) effect than *Nfkbia*-targeting ASOs (**Figure 6G-I**). However, the fact that the mathematical model reproduces the experimentally observed change in AUC by adjusting only *Nfkbia* as observed experimentally suggests that the dynamic changes observed are largely due to specific targeting of IκBa.

Overall, we have predicted through mathematical modeling that small changes between clone B and clone R in the expression levels of the negative feedback can shift the dynamics from a sharp versus a persistent NF-κB response. Our experiments with ASOs show how even a mild targeted modulation of the expression of IκBα can alter the dynamics of clone R to resemble that of clone B, and therefore to reprogram it from a sharp (and oscillatory) NF-κB activation to a persistent response, as predicted by our mathematical models.

## DISCUSSION

### Clones from a population of MEFs with heterogeneous NF-κB dynamics have distinct NF-κB dynamics

Single-cell imaging studies of NF-κB dynamics have shown that there is a high degree of heterogeneity within homogeneous cell populations (Lee et al., 2014; Nelson et al., 2004; Paszek et al., 2010; Sung et al., 2009; Tay et al., 2010; Q. Zhang et al., 2017). We find that clonal populations from a cell line of fibroblasts derived from a single mouse embryo have distinct NF-κB dynamics, similar to those observed in single cells of the original population. This indicates that NF-κB dynamic heterogeneity might be due in part to the coexistence of sub-populations of cells that do respond distinctly to the stimuli. Extrinsic factors such as the cell cycle phase have been shown to affect NF-κB dynamics (Ankers et al., 2016) but we cannot attribute inter-clonal differences to differences in the cell cycle. Instead, we show here that the expression levels of genes coding for key elements of the NF-κB signaling pathway can explain the clonal differences.

### Transcriptomics is predictive of the clonal response to stimuli

The link between NF-κB dynamics and transcription is typically analyzed in one direction: how does NF-κB nuclear localization dynamics drive gene expression? (Lane et al., 2017; Lee et al., 2014; Sen et al., 2020; Tay et al., 2010; Zambrano et al., 2016). Here we consider the same question in the other direction: how can transcriptomic differences affect dynamics? We developed a computational framework based on transcriptomic-constrained estimation of the number of activating complexes formed upon interaction of the receptors with different ligands, which in turn has been shown to correlate with activation of the IKK complex and NF-κB activation (Cruz et al., 2021). Modeling suggested that the limiting factor in the activation of NF-κB in response to TNF-α and IL-1*β* is the abundance of the cognate receptor; the abundance of signal transducers is in large excess relative to receptors and hence is not the limiting factor in NF-κB activation. Our transcriptome-constrained estimations indicated that the ranking of clone responsiveness to IL-1*β* and TNF-α would be different, and experimental evidence confirmed the prediction. Notably, the notion that the abundance of receptor transcripts can be used to predict responsiveness also holds for other systems (Su et al., 2022).

We also show here how differences upstream of NF-κB, combined with the expression levels of IκBa, which exerts the main negative feedback on NF-κB, predict whether the NF-κB response will be sharp or persistent. Furthermore, the experimental data show that a moderate reduction in IκBα level induced by antisense oligonucleotides can lead to relatively large differences in the NF-κB dynamics. Consistently, the knock-out of the gene *Nfkbia* has been shown to induce a change from oscillatory to sustained dynamics (Cheng et al., 2021) (Adelaja et al., 2021). We speculate that incorporating further transcriptional differences in the negative feedbacks would allow us to reproduce other differences between the clones, such as the level of heterogeneity (Paszek et al., 2010) or differences in the responses to subsequent stimuli (Adamson et al., 2016).

Part of the success of our approach is probably due to the biological homogeneity of the clones, which derive from cells of the same type from a single embryo. We assume that differences in the transcriptional levels between our cell populations are informative of differences in the protein levels; while this is not necessarily true for any two cell populations, we expect it to hold true for clonal populations derived from cell lines. We hypothesize that a similar analytical approach could be used to predict the relative difference in the dynamic response of other TFs in other relatively homogeneous cell populations, as for example in cells derived from the same tissue or in clones within a tumor mass (Greaves and Maley, 2012) that might respond differently to therapy (Paek et al., 2016).

### NF-κB oscillatory phenotype is fuzzy

NF-κB nuclear localization dynamics in single living cells was observed for the first time more than 15 years ago (Nelson et al., 2004) and since then its oscillatory nature was subject to discussion (Barken et al., 2005; Nelson et al., 2005). More recently, we argued that NF-κB displays damped oscillations (Zambrano et al., 2016) as compared to the sustained oscillations reported by others (Kellogg and Tay, 2015). Our present work shows that classification of NF-κB dynamics is not necessarily binary. Within a cell population, we can generate clonal populations that are more and less prone to oscillate. For circadian oscillators as well it was found that clonal populations have different oscillatory features (Li et al., 2020). For our cells, we show that this largely depends on the expression level of genes belonging to the NF-κB regulatory circuit. If small transcriptional differences can affect the dynamics of NF-κB, it is not surprising that different cell types have completely different oscillatory phenotypes.

### Stimulus specificity in the NF-κB response is clone-dependent

The increasing availability of single-cell data on NF-κB dynamics has shown that it is possible for cells to discriminate between stimulus type (Adelaja et al., 2021; Cheng et al., 2021; Martin et al., 2020), dose (Tay et al., 2010; Zambrano et al., 2014a; Q. Zhang et al., 2017) and dynamic profile (Ashall et al., 2009; Lee et al., 2016; Zambrano et al., 2016). Here, we find that two MEF clones from the same embryo, clone G and clone B, do respond differently to two different stimuli: clone G responds strongly to IL-1*β*, while clone B responds strongly to both TNF-α and IL-1*β*. On the other hand, clone R produces a sharper and more oscillatory NF-κB response than clone B. This indicates that stimulus specificity in NF-κB dynamics is clone-dependent and, all the more so, it will vary among cell types within the same organism. This is in line with a recent study where a synthetic version of the NF-κB system ectopically expressed in yeast (Zhang et al., 2017) displays different types of dynamics and responses by manipulating the expression of key genes within the genetic circuit.Furthermore, since immortalized MEFs maintain certain characteristics of primary MEFs (Beg and Baltimore, 1996), our work suggests that a similar fine-tuning can take place naturally in primary mammalian cells; although a recently developed reporter mouse (Rahman et al., 2022) has allowed to show that primary cells and cell lines can have different NF-κB dynamical features. Only further analysis of primary cells will allow us to address this question.

### Origin of the transcriptional differences across clones

We show that the dynamical differences observed between clones are robust and persist over time and cell culturing. Our bioinformatic analysis could not detect any variant among clones in the coding sequence of genes involved in the NF-κB response. We inferred a substitution rate in the genome of the clones in the order of 5·10^−7^ per base pair, which would translate to <100 substitutions per haploid genome. This substitution rate is consistent with somatic variability of cells from the same organism (Amand et al., 2016; Milholland et al., 2017). However RNA-sequencing based analysis cannot detect other genetic changes, such as sequence changes in the regulatory regions or copy number variations. These could well lead to the small (although robust) differences in gene expression reported here. The other possibility is epigenomic variation between different cells, i.e. in DNA methylation or chromatin accessibility of gene control elements. We find that the most visible difference in the transcriptomics of the different clonal populations is related to developmental programs, which indeed involve epigenetic variations. Whether expression of different developmental programs can give rise to the difference in expression of the genes that affect the NF-κB dynamic response remains to be proved, but we speculate that this is at least likely.

### Cell to cell differences within clones

We find that the NF-κB dynamic response of the cells within each clonal population is still heterogeneous. We speculate that this cell-to-cell variability can have a purely stochastic component related to the probabilistic nature of the activation of gene transcription. Indeed, we recently found experimentally that even highly transcribed genes under the control of NF-κB like the one encoding for IκBα are transcribed stochastically (Zambrano et al., 2020). Thus, the same cell might be oscillatory or not at different times, depending on how recently it had a burst of IκBα transcription and translation. Future studies will be needed to connect the transcriptional history of each single cell with its NF-κB dynamics.

In sum, our work shows that clonal populations from a homogeneous cell line have distinct NF-κB dynamic in response to stimuli, due to small (less than twofold) differences in the expression levels of genes belonging to the NF-κB signaling pathway. This indicates that sub-populations with heritably distinct NF-κB dynamics coexist in the original cell population. However, some heterogeneity remains between cells of the same clone, which we suggest might be due to noisy transcriptional bursts in individual cells. We speculate that analogous mechanisms might also diversify the dynamic response of other TFs to external cues.

## ACKNOWLEDGEMENTS

We acknowledge Prof. Manolis Pasparakis for sharing with us the immortalized MEF population from a single embryo, Prof. Roberto Sitia for suggesting the initial experiment and Davide Mazza for critical revision of the manuscript. We received funding from Ospedale San Raffaele (OSR Seed Grant to SZ), San Raffaele University (predoctoral fellowship to CK) and Associazione Italiana Ricerca sul Cancro (AIRC 2017-ID.18687, project PI AA). Part of the present work was performed by CK in partial fulfillment of the requirements for obtaining the PhD degree at Vita-Salute San Raffaele University, Milano, Italy.

## AUTHOR CONTRIBUTION

Conceptualization: SZ. Supervision: SZ, MEB, AA. Visualization: SZ, CK, EM, MEB. Formal Analysis: CK, SZ, EM, AA. Investigation: CK, SZ, EM, JGM, FB. Writing-Original draft: SZ, CK, MEB Writing-review and Editing: SZ, CK, EM, MEB, AA, JGM, FB. Funding acquisition: SZ, MEB, AA.

## FIGURE LEGENDS

## RESOURCE AVAILABILITY

### Lead Contact

Requests for resources and reagents should be directed to the lead contact, Samuel Zambrano.

## Materials availability

All unique reagents generated in this study are available, with a completed Material Transfer Agreement, from the lead contact upon request.

## Data and code availability

- Data on NF-κB dynamics for single cells allowing to reproduce the analysis Figures 1, 2, and 5 are provided in the Supplementary Data File. Gene expression level for the conditions and biological replicates described in Figure 3, 4 and 5 are provided as .csv files and in excel format in the Supplementary Data File.
- Imaging routines to extract NF-κB signaling dynamics in single cells from time lapse movies and mathematical modeling-related code (transcriptomic-constrained estimations of number of complexes and signaling dynamics) can be downloaded at https://github.com/SZambranoS/. Bioinformatics analysis tools are available as described in the method details section.
- Any additional information required to reanalyze the data reported in this paper is available from the lead contact upon request.

## EXPERIMENTAL MODEL

### Cell line and cell culture

GFP-p65 knock-in mouse embryonic fibroblasts (MEFs) were kindly provided by M. Pasparakis. MEFs were derived from a single male embryo of a homozygous knock-in GFP-p65 expressing mouse model, using standard protocols (De Lorenzi et al., 2009) and immortalized by serial passaging following the “3T3 method” that typically involves 20-30 passages (Amand et al., 2016). Clones were generated from a freshly frozen vial of these MEFs, and aliquots of each clone were frozen and then used for few passages, to keep at a minimum the number of passages. The cells were cultured in phenol-red free DMEM supplemented with 10% FCS, 50 mM b-mercaptoethanol, 1x L-glutamine, 1x pen/strep, 1x sodium pyruvate and 1x non-essential amino acids. MEFs were subcultured every 2-3 days before they reached 100% confluency and kept at 37°C and 5% CO2.

### Generation of the clonal populations by single cell cloning

MEFs were harvested by 1x Trypsin solution and counted. Final concentration of 5 cells/ml was achieved by serial dilutions and 100 μl of the cell suspension per well were pipetted to a 96-well plate. The plate was screened for single colonies and selected colonies were then expanded.

## METHOD DETAILS

### Cell treatments

Where indicated the cells were stimulated with the final concentration of 10 ng/ml of recombinant human TNF-α protein (R&D Systems) or 100 ng/ml of recombinant human interleukin 1 beta (IL-1*β*, PeproTech).

### Live cell imaging

Live cell imaging of GFP-p65 knock-in MEFs was performed as in (Zambrano et al., 2016). We used a Leica TCS SP5 confocal microscope with an incubation system where cells were stably maintained at 37°C in 5% CO2. Time-lapse images were acquired at 6 min intervals for up to 10 hr. We used a low magnification objective (20x, 0.5 NA) and an open pinhole (Airy 3), ensuring that the image depth (10.7 μm) contains the thickness of the whole cell so that images capture the total cell fluorescence. GFP-p65 is imaged with the 488 nm Argon laser (GFP channel) while Hoechst 33342 stained nuclei are imaged with the low energy 405 nm UV diode laser at 5% of its maximum intensity (HOE channel). The staining was performed at room temperature for 10-15 minutes using NucBlue™ (Live ReadyProbes™ Reagent, ThermoFischer), 1:100 v/v and incubated 10-15 min at RT. We showed previously in this same cell line that imaging and staining do not interfere with the response to TNF-a, the cell’s viability and ability to replicate (Zambrano et al., 2016), so we exclude any relevant phototoxicity effects of our imaging. Images were acquired as 16 bit, 1024×1024, TIFF files. Experiment replicates were performed on different days. In each experiment we typically imaged more than one clone in different wells of an 8-well labtek.

### Automated quantification of NF-κB dynamics in single living cells

To quantify NF-κB nuclear localization dynamics in living cells, we follow our previously described procedure of normalizing the average nuclear signal intensity by the average cytosolic fluorescence intensity (Zambrano et al., 2016) to obtain the nuclear to cytosolic intensity (NCI), also used by others (Kellogg and Tay, 2015; Paszek et al., 2010). We improved our custom-made routines that run on Matlab R2015 and are made available online. In short, nuclei are segmented based on the intensity of the HOE channel, and nuclear masks are used to compute the nuclear average NF-κB intensity in each cell. In order to estimate the average cytoplasmic NF-κB intensity, first the background was computed by dividing the image in small 32-pixel tiles and using the one with the smallest average intensity as a background estimation.. After this, pixels belonging to the cytoplasm are those with intensity above the background on a ring around each nucleus of width 0.5 times the nuclear radius. Tracking of cells between frames is performed through an optimized algorithm based on the Hungarian linker method (Careccia, 2019). Cells are discarded upon abrupt changes of the nuclear and/or cytosolic areas, indicative of erroneous tracking or cell death or mitosis. The resulting NCI profiles, that we refer to as NF-κB dynamic profiles, where smoothened using the Matlab function *smooth*.

### Stochastic clustering of the NF-κB dynamic profiles

We performed an unsupervised clustering of NF-κB dynamic profiles from cells of the 8 clones and the pool using the k-means algorithm (k=9) implemented on Matlab and based on the euclidean distance between profiles. Since we have hundreds of cells per clone, in each realization we randomly picked 50 profiles from each population. The profiles are clustered then in 9 groups (**Figure S1B**), and we compute the number of cells from each clonal population in each cluster (**Figure S1C**). In each realization we compute *p*_*ik*_, representing the fraction of trajectories of clone *k* that are present in the cluster where the clone *i* has a higher number of clustered profiles. For *k=i*, it represents the fraction of cells of the clone *i* in the cluster where it is more represented. The result for a single realization is shown in **Figure S1D**, the average of many realizations is on **Figure 1F**. For 500 realizations we computed the disorder parameter defined as 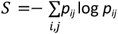. For each of them we repeated the procedure but randomly assigning the selected profiles to the 9 populations, and calculating for each clustering the disorder parameter. The disorder parameter was always higher for cells randomly assigned to the populations, indicating further that NF-κB dynamic profiles are clustered prevalently according to the clonal population of origin.

### Analysis of NF-κB dynamics

To extract the dynamic features of the NCI time series we followed the same procedure as in (Zambrano et al., 2014, 2016). In short, NCI series are smoothed and peaks are detected using standard Matlab functions (*smooth* and *findpeaks*, respectively) and those with a prominence θ>0.15 are considered real peaks. This value is well beyond the prominence of noisy peaks found in this type of datasets (Zambrano et al., 2016) and provides a reasonably good compromise between the need to ignore noise peaks and the need to detect small peaks of valuable dynamical information (e.g. to classify an NF-κB dynamic profile as oscillating or not). The timing of the peak was determined considering the maximum value. Instead, the area under the curve is calculated as the integral in the time interval considered of NCI(t).

### Cell cycle analysis

Cell cycle analysis was done as described in (Brambilla et al., 2020): MEFs were harvested and fixed with cold 70% ethanol and kept overnight at -20°C. Cells were then washed once with 5% FBS/PBS and stained with PBS containing 10 μg/ml propidium iodide and 10 μg/ml RNAse A for 1 hour at room temperature. Samples were then read at a cytometer using a 488 nm laser.

### Cell cycle computational correction

Clone B has a stronger response than clone R, and it has a higher fraction of cells in the S-phase. This could be the source of the stronger response in clone B so, to computationally correct for this, we generated an artificial cell-cycle corrected dataset of NCI time series of clone R where the time series where sorted by their response and the top 25% of the population was assumed to be the S-phase high responders (Ankers et al., 2016). To increase their percentage to 43% (as clone R) we discarded 42% of the profiles, those of lower responses, to “match” the fraction of cells in S-phase in clone B (see Table 1). The resulting dataset has a higher AUC value than the original but still a lower value of the AUC than clone B (**Figure S2J**) which makes it unlikely that cell cycle is the key driver of this inter-clonal difference. Furthermore, clones B and G have very similar cell cycle distribution and even so their dynamics upon TNF-α are markedly different, as detailed in the main text.

### RNA isolation and real time PCR

1.5×10^5^ MEFs were plated on a 6-well plate a day before the extraction. Total RNA was isolated using the NucleoSpin RNA II kit (Macherey-Nagel). The amount of RNA was determined using a NanoDrop spectrophotometer (Thermo Fisher Scientific) and 1 μg was then reverse transcribed using SuperScript IV Reverse Transcriptase (Thermo Fisher Scientific). qPCR was performed using LightCycler 480 SYBR Green I Master (Roche). The expression of *I*κ*Ba* was checked using following primers:

*I*κ*Ba* forward: 5’ CTTGGCTGTGATCACCAACCAG 3’

*I*κ*Ba* reverse: 5’ CGAAACCAGGTCAGGATTCTGC 3’

### RNA sequencing and Bioinformatic analysis

Libraries for Illumina NGS were prepared as described in (Brambilla et al., 2020). After trimming the adapter sequences (cutadapt, https://cutadapt.readthedocs.io) reads were mapped to mouse genome (mm10) using hisat2 (http://daehwankimlab.github.io/hisat2/) using parameters “-p 20 -5 5”.

Read counting was performed using featureCounts from the Subread Package and features displaying less than 10 reads were filtered out. Differential expression analysis was performed using DESeq2 (https://bioconductor.org/packages/release/bioc/html/DESeq2.html) with the following design formula “=∼ 1 + clone + treatment + clone:treatment”.

Principal component analysis (PCA) was performed using Matlab, keeping only genes for which RPKM>1 in at least five samples. Additional RNA-seq data from several mouse tissues were retrieved from the ENCODE Database (https://www.encodeproject.org). Volcano plots were generated using Matlab and p-values derived from the t-test statistics. Gene ontology was performed using the “*clusterProfiler*” R package (https://bioconductor.org/packages/release/bioc/html/clusterProfiler.html). Heatmap and hierarchical clustering was performed using the “*Pheatmap*” R package.

Genes annotated in the mouse TNF-signaling were retrieved from WikiPathway (https://www.wikipathways.org/index.php/Pathway:WP246)

ssGSEA analysis: Genes from the “TNF-α to NF-κB” and “IL-1*β* to NF-κB” lists were ranked according to their log normalized expression (log2 FPKM + pseudocount) and for each biological replicate (n=5), normalized enrichment scores (NES) were calculated (Barbie et al., 2009). The NES reflects the degree to which the gene list is coordinately up or down-regulated within each clone.

### Estimation of the genetic differences between clonal populations

The SNP calling step was performed using the GATK 3.6 toolkit (McKenna et al., 2010) in order to split splice junction reads, to recalibrate quality scores and to call variants. To minimize false positive variants the GATK Variant filtration tool was used using the following parameters:

“--filterExpression QD < 5.0 --filterExpression DP < 10 --filterExpression ReadPosRankSum < -8.0 --filterExpression MQRankSum < -12.5 --filterExpression MQ < 40.0 --filterExpression FS > 60.0”. Nucleotide positions with heterozygosity scores < 0.10 were excluded as previously described (Adetunji et al., 2019).

We called SNPs with different levels of confidence based on three different coverage cut-offs (5x, 10x, 20x) and then we calculated the number of unique SNPs for each clone. We find that our clones differ in a range of 200-400 nucleotides (**Table 2**) which, once divided by the length of the genome at the specific coverage cut-off, provided us a “mutation rate” of approximately 5·10^−7^ per base pair (**Figure S3B**). Interestingly, this is of the order of magnitude of the somatic mutation rate found between somatic cells from the same mouse (Milholland et al., 2017) indicating how the frequency of SNPs for our clones correspond to “somatic differences” that can be found between cells of the same organism. Of note, our cells come from the same embryo and immortalization by serial passages requires a few dozen cell replications (Amand et al., 2016; Todaro and Green, 1963). Moreover, no mutations on NF-κB-related genes of the *wikipathways* database were identified.

**Table 2:** Length of the genome covered for different coverage thresholds (in bp) using RNA-seq reads, and number of different SNPs identified between clones.

| coverage\length | genome covered R | genome covered B | genome covered G | Diff. B vs G | Diff. B vs R | Diff. G vs B |
| --- | --- | --- | --- | --- | --- | --- |
| 5x | 560805040 | 614001397 | 576431940 | 351 | 301 | 338 |
| 10x | 470022271 | 513879107 | 480558887 | 193 | 172 | 179 |
| 20x | 384083108 | 425738181 | 392300809 | 163 | 108 | 135 |

**Table 3:**
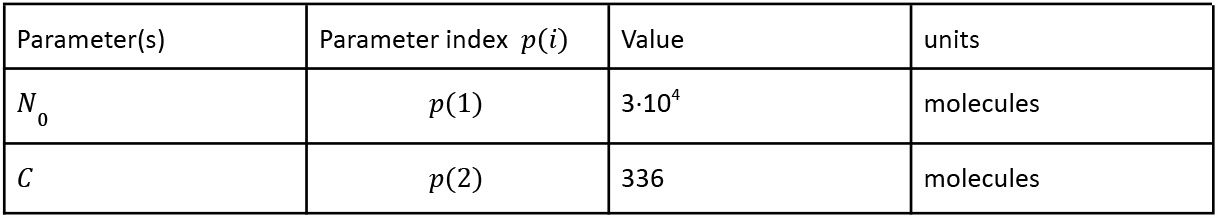
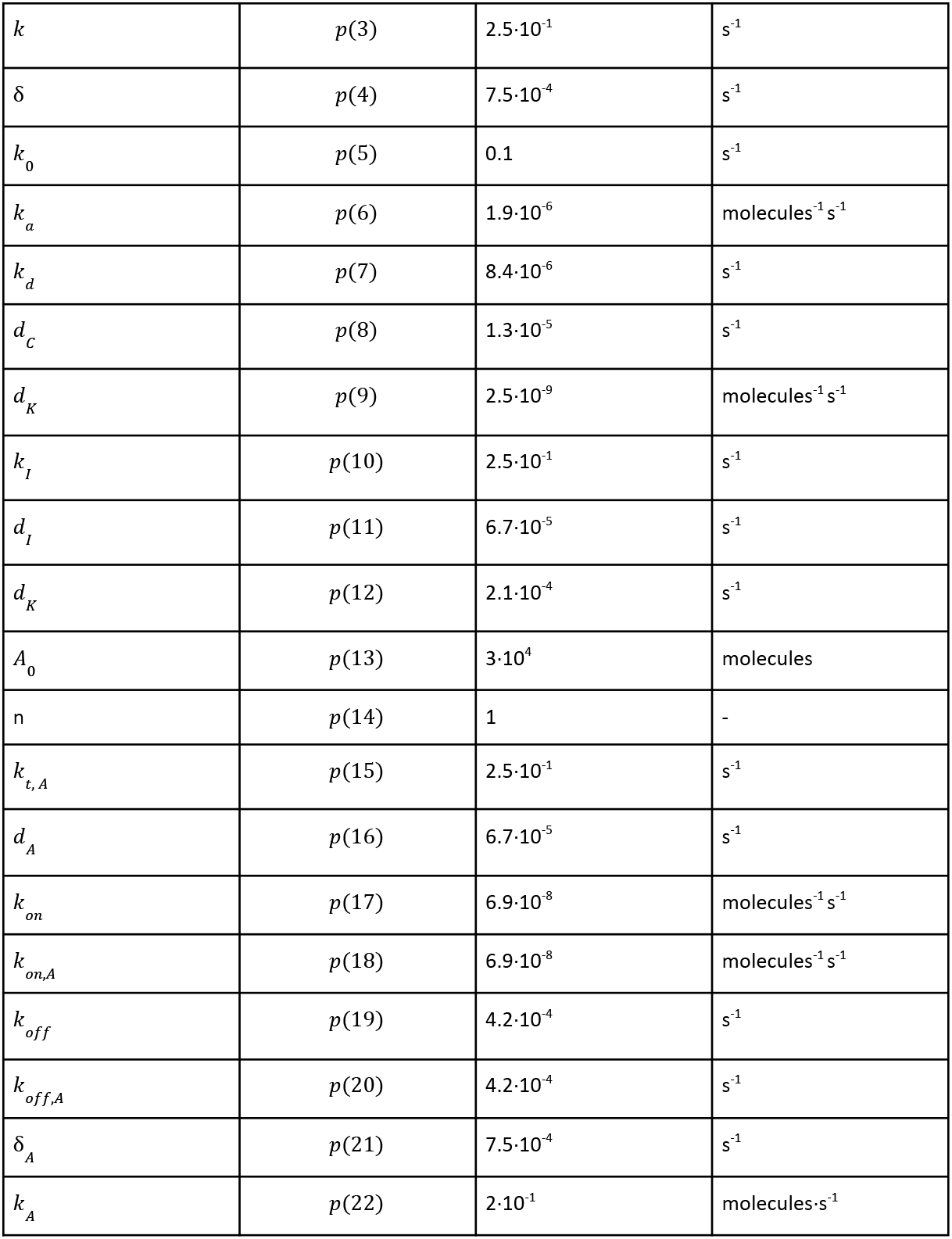
Parameters of the mathematical model: Starting parameters considered in all our simulations.

### Assumption of proportionality between RNA and protein levels among clones

This important assumption in our work relies on the following theoretical consideration. Proteins can be considered to be translated from mRNAs at a constant rate *k* and degraded with a constant rates γ. In the equilibrium,we have that the levels of the protein (*p*) and of mRNA (*m*) are related by the equation: 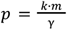(Suter et al., 2011). Since our clones derive from a relatively homogeneous population, we assume that our clones have similar translation and degradation rates for each protein, and hence the proportions between RNA and protein levels shall be similar 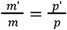, as for p65 (**Figure S4A**). Moderate deviations from this rule would give qualitatively similar results in our simulations.

### Complex formation simulations and estimation of the equilibrium concentrations

To estimate the differences between clones in the number of activating complexes formed after interaction between ligand and the receptor we describe their formation as a sequence of association-dissociation reactions. For the sake of simplicity, and considering also experimental data on the dynamics of complex formation (Cruz et al., 2021), we consider that those reactions take place very quickly compared with NF-κB response, so their equilibrium concentration is the only relevant magnitude to determine the short-term NF-κB response.

To evaluate the equilibrium concentrations, we make the following considerations. Each reaction of the formation of a complex (*A*: *B*) of biochemical species *A* and *B* can be written as:

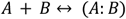

While the evolution in time of the number of copies of each species, that we represent with the same letters, is given by

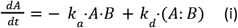

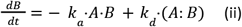

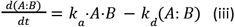

Where *k*_*a*_ and *k*_*d*_ are the association and dissociation rates, respectively, of the species to form the complex. If we denote by *A*_0_ and *B*_0_ the total amount of the biochemical species *A* and *B* (the sum of the free and the complex-forming ones), the equilibrium concentrations *A*_*eq*_, *B*_*eq*_ and (*A*: *B*)_*eq*_satisfy the equations:

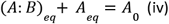

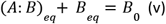

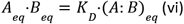

Where *K* _*D*_ = *k* _*d*_ /*k* _*a*_ is the dissociation constant. If we specify the species amounts in number of molecules, *K*_*D*_ is also given in numbers of molecules (when specified in concentrations, the dissociation constant has units of concentration).

It is relatively easy to see that the equilibrium concentrations can be found solving a second-degree equation. In **Figure S4D** we show simulations of this system in which we consider that *A*(0) = *A*_0_, *B*(0) = *B*_0_ and hence (*A*: *B*)(0) = 0, showing how they all converge to the equilibrium values computed solving the ODEs (i)-(iii) or finding the zeros of (iv)-(vi). The panels in **Figure S4D**, from left to right, show the evolution of the species towards equilibrium in a situation where *B*_0_ < *A*_0_ : if *K*_*D*_ = 0, then (*A*: *B*) _*eq*_ = *B*_0_ (the smaller abundance) ; if *K* _*D*_ ≪*B* _0_ then (*A*: *B*) _*eq*_ ≃ *B* _0_ ; if *B* _0_ ≤ *K* _*D*_≤ *A* _0_ then also *B*_0_ ≤ (*A*: *B*) _*eq*_ ≤ *A* _0_ and finally if *K* _*D*_ ≫ *A* _0_, (*A*: *B*) _*eq*_ ≃ 0. Reasoning along these lines, it is easy to see that in the process of association of many species to form a complex, the upper bound of the number of complexes is given by the less abundant species.

### Transcriptomic-constrained estimation on the number of activating complexes in the clones upon stimuli

The formation of the Complex I upon TNF-α stimulation (DeFelice et al., 2019; Hayden and Ghosh, 2008; Hsu et al., 1996; Lee et al., 2016; Wilson et al., 2009) or what we call here the “activating complex for TNF-α”, can be represented as the following biochemical reactions:

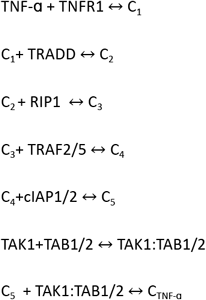

Notice that we denote the complex emerging from the addition of a new element as C_n_ for the sake of simplicity. The final activating complex is C_TNF-a_. The list of the genes coding for the proteins involved in the reactions, our “TNF-α to NF-κB” list, is : *‘Tnfrsf1a’, ‘Tradd’, ‘Ripk1’, ‘Traf2’, ‘Traf5’, ‘Birc2’, ‘Birc3’, ‘Map3k7’, ‘Tab1’, ‘Tab2’*.

Analogously, the activation of NF-κB upon IL-1*β* follows the formation of its own activating complex upon the interaction of the dimeric receptor with its ligand: following the reactions (Martin and Wesche, 2002; Rhyasen and Starczynowski, 2015):

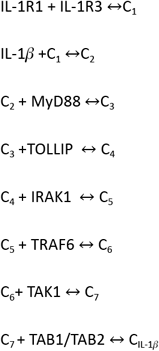

The list of the genes coding for the proteins involved in the reactions, our “IL-1*β* to NF-κB” list, is: ‘*Il1r1*’;’*Il1rap*’;’*Myd88*’;’*Irak1*’;’*Map3k7*’;’*Traf6*’;’*Tab1*’;’*Tab2*’. The activating complex is denoted by C_IL-1*β*_.

The description above is a simplification where we have deliberately neglected the precise stoichiometry of each reaction (which in some cases is not so clear); however, we also implemented simulations using other stoichiometries postulated in the literature with analogous results (reproducible with the software provided).

In principle, if we knew the copy number of each protein and the dissociation constant of each of the association-dissociation reactions above, we shall be able to determine the abundances of the complexes C_TNF-a_ and C_IL-1*β*_ in equilibrium (which, as we said, we assume to happen fast). However, we do not know them, but we can make simulations to compute how their numbers compare between clones under reasonable assumptions:

- First, as described previously, we assume that clonal populations have similar translation rates and protein degradations for each protein, so the ratios between the amount of transcript and protein of each species should be similar. Since we have 5 replicates of each condition, we consider that for each simulation we have 5 values of the relative protein abundance for each protein involved. This also provides us with an experimental error that will contribute to the uncertainty of our estimates.
- Second, we know that there is a difference in the order of magnitude of the number of copies between certain proteins in these reactions and the remaining ones; in particular the protein copies of TNFR1 are about Δ≳3 orders of magnitude less abundant than the other proteins leading to C_TNF-a_ (O(10^3^) vs O(10^6^)), and IL-1R proteins are about Δ≳3 orders of magnitude less abundant than the remaining proteins leading to the complex C_IL-1*β*_ (O(10^2^ - 10^3^ vs O(10^6^)) (Cruz et al., 2021; Hwang et al., 2015). In our cells *Il1rap* is much less expressed than *Il1r1*, we consider the ratios of protein abundance to match the RNA expression for these two highly homologous proteins. These receptor proteins will act as limiting factors in the abundance of the complexes.

Under these two assumptions, we can make estimates of the activating complex abundances C_IL-1*β*_ and C_TNF-a_ for each clonal population by randomizing the copy numbers of each protein species, but preserving the ratios between expression in the clones specified by transcriptomic data and the difference in order of magnitudes Δ between the limiting factors (the receptors) and the remaining proteins of the cascade.

Specifically, simulations were performed by stochastically assigning abundances of the proteins involved in the complexes keeping the experimentally determined Δ values and by using different *K*_*D*_ values between 1 and the maximum value considered, which was the maximum protein abundance, 10^6^ copies. Once randomized, we calculated deterministically the equilibrium concentrations C_n_ in a sequential way, using the same numerical tools described in the previous section, up to C_TNF-a_ and C_IL-1*β*_. The ratios of the resulting number of activating complexes between clones are then informative of the expected FC difference in copy numbers of the activating complex between each of them. These values will be different in each simulation of these transcriptomic-constrained simulations, but we can find their mean value and its standard deviation for given *K*_*D*_ and Δ values.

It is worth discussing two limit cases. First, for large Δ values in principle the ratios between the numbers C_TNF-a_ and C_IL-1*β*_ will be dominated by the ratios of the less abundant proteins in the simulations, which in turn will be proportional to the ratios of the RNA expression. We indeed find that this convergence takes place really fast for values of Δ much smaller than the experimentally estimated ones if the values of *K*_*D*_ are moderate (e.g. **Figures 4E** and **S4E, Figures 5D** and **S5D**). Second, for large *K*_*D*_ values (comparable to the copy number of the proteins considered here (O (10^6^)) we might expect that some step of the complex formation reactions will lead to a very small number of resulting complexes (consider the simple two species case of **Figure S4D**). This results in a high variability in the number of complexes estimated for large *K*_*D*_ as we observe in **Figure S4F** and **Figure S5F**. Those panels show how such variability is mitigated for sufficiently large Δ by the mathematical reasoning sketched above.

### Mathematical model of NF-κB signaling

We use here a slightly modified version of our model of NF-κB dynamics (Zambrano et al., 2014b; Zambrano et al., 2016), that we briefly describe below.

As described in different works and experimentally observed, NF-κB activation follows the formation of a number of activating complexes *C* downstream the ligand-bound receptor, where the kinase complex IKK is activated proportionally to the amount of activating complex formed *C* upon stimuli (C_TNF-a_ or C_IL-1*β*_) through the constant *k* _0_. Hence the evolution of the abundance of the active IKK complex, that we denote *K*, can be written as:

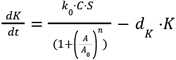

The amount of activating complex is regulated by the amount of the protein A20, that we denote *A*, through a hill function of parameters *A*_0_ and *n*, and will also depend on the inactivation rate of the kinase complex *d*_*K*_ and the presence (*S* = 1) or absence (*S* = 0) of the external inflammatory signal.

Importantly, this simple model of kinase activation reproduces the linear relationship between number of complexes *C* and NF-κB response (measured as maximum of the nuclear to cytosolic NF-κB intensity, **Figure 4G**), in agreement with what has been observed experimentally (Cruz et al., 2021).

In this new model, the amount of free nuclear NF-κB, *N*, depends on its continuous association-dissociation with the IκBα inhibitor protein, whose abundance we represent as *I*, to form the complex (cytosolic) form (*N*: *I*). The active kinase can degrade the inhibitor in the complex, so:

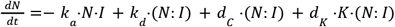

Where *k*_*a*_ and *k*_*d*_ are the association and dissociation rates, respectively, *d*_*K*_ is the degradation rate of the inhibitor in (*N*: *I*) due to the presence of the active kinase, while *d*_*C*_ is the spontaneous degradation of the inhibitor in (*N*: *I*). The evolution of the complex abundance is given by:

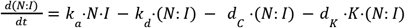

So the total amount of NF-κB (free plus bound) is constant *N* + (*N*: *I*) = *N*_0_.

For the free inhibitor, beyond the association and dissociation with NF-κB, the evolution will depend also on the translation rate *k*_*I*_ protein degradation rate *d*_*I*_ of the available mRNA, that we denote as *R*, and on the inhibitor’s own so:

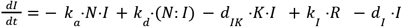

The mRNA expression level of the mRNA amount *R* is summarized in the following equations that model the first main negative feedback of the system IκBα, determined by the level of activity of the gene *G* coding for this inhibitor:

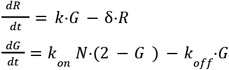

where δ is the degradation rate of the mRNA and *k* is the transcription rate, while and *k*_*on*_ and *k*_*off*_ are the gene activation-inactivation rates respectively.

The amount of A20, the second negative feedback of the system that operates upstream and we denote as *A*, is governed by an analogous set of equations that encode for the time evolution of its protein abundance, *A*, its available mRNA, that we denote as *R*_*A*_, and on the activity of the gene encoding for it, that we denote *G*_*A*_ :

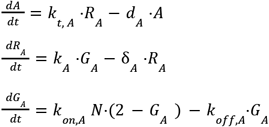

With its own parameters governing gene activation/inactivation, transcription and translation kinetics, in an analogous way to the parameters of the IκBα feedback. The starting parameters of the model are specified in Table 3. Finally, the total amount of NF-κB is referred to as *N*_0_. We denote as *p* the vector of parameters of the system.

*N*(*p, t*) is the evolution of time of the nuclear amount *N* obtained using our model and a given parameter set, and the nuclear to cytosolic ratio *NCI*(*p, t*) is calculated as

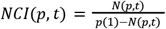

Notice that, as specified by table 3, *p*(1) is the total amount of NF-κB.

In our simulations, the model is integrated typically for up to 4 hours. We define the maximum value as

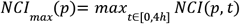

and the area under the curve as

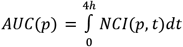

Finally, if *R*(*p, t*) represents the RNA of *Nfkbia* in a given simulation for parameters *p*, we define its peak value as:

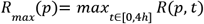

### Experimentally constrained mathematical modeling of NF-κB dynamics

In the paper we perform numerical simulations constrained by our experimental data, to assess in general to what extent small experimental differences between clones B, R and G can explain the different dynamic responses that we observe. To this aim, we use random sets of parameters of our model that and then we adjust only some of them (to match certain experimental observations), and use the model to see if this parameter adjustment results in changes in the NF-κB dynamics compatible with those observed experimentally. This operation is repeated starting from different randomized parameter sets that are “prototypical randomizations of clone R” that are taken as a starting point, on which we apply the experimentally-derived corrections of the parameters described below and then analyze the effect in the dynamics. We describe the procedure in further detail below.

### Randomization of parameters to obtain prototypical NF-κB dynamics of clone R

To generate prototypical parameters of clone *R, p*_*R*_, the parameters of Table 3, noted *p*_0_, are randomized up to 2-fold (multiplying each of them by a different factor 2^ξ^ with ξ a random number in the [-1,1] interval); a randomized parameter set *p*_*R*_ is considered prototypical if the maximum value of *NCI*(*p*_*R*_, *t*) is between 1.5 and 3.5, as experimentally observed by us and others upon TNF-α.

### Simulating the effect of small transcriptional differences in the dynamics

#### Reproducing NF-κB amount differences among the clones

A first simple example is that of **Figure S4G** where we perform a simulation with the parameters specified in the Table 3 to obtain a simulated trajectory of clone R (red line), so in this case *p*_*R*_ =*p*_0_. Then we define parameter sets for simulations of clone B and clone G by changing the NF-κB amount in our model (*N*_0_) with respect to that of R, following the experimentally observed ratios of B vs R and G vs R (**Figure S4A**). Mathematically,

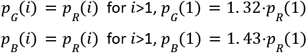

**Figure S4G** shows *NCI*(*p*_*R*_, *t*), *NCI*(*p* _*G*_, *t*) and *NCI*(*p* _*B*_, *t*), The simulation shows clearly that changing NF-κB amount does not reproduce the experimental observed differences in the NF-κB response between clones. Analogously, to estimate the effect of just varying NF-κB amount in general, as shown in **Figure 6E**, we generate randomized parameters *p* _*R*_ and increase the amount of NF-κB as specified above to obtain *p* _*B*_ and then calculate the fold change in the AUC; we find that it is a relatively small fold change in all cases compared to the experimental observations.

#### Modulating NF-κB amount and estimating the FC in the amount of activating complexes

**Figure 4G** shows that a variation of the activating complexes can lead to a variation in the response predicted by the model, as observed experimentally (Cruz et al., 2021) (inset of **Figure 4G**). **Figure S4H** shows simulations of the change in the response (taking as reference value clone R) for different changes in the amount of activating complexes (*C*) and the amount of NF-κB in the model (*N* _0_). This allows us to estimate the FC variation in the number of activating complexes and in the amount of NF-κB of B respect to R and of G respect to R that reproduce the observed changes in the NF-κB response upon TNF-α and IL-1*β*. We repeated the operation using parameter randomizations to obtain different prototypical parameters of R, *p* _*R*_, as described above, and for each of them generate new parameters sets that differ only in the NF-κB amount and in the amount of activating complex, that we fit to reproduce the difference in the responses, i.e.:

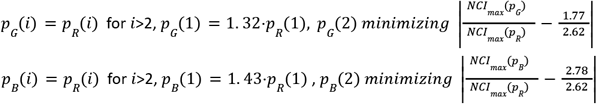

Notice that each resulting parameter sets will differ from *p* in two parameter values. Hence *p* _*G*_(2) and *p* _*B*_(2) are the result of minimizing indicated cost functions that have a minimum when the ratio of the maximum of *NCI*(*p*) *max* numerically computed for each parameter set (*p* _*B*_ and *p* _*G*_) and *NCI max* (*p* _*R*_) matches the experimental differences between the averages of the maximum response of the clones considered. The values are found through Matlab optimization tool fminbnd. Hence, the model estimates of the FC variation of the number of activating complexes *C* that reproduces the experimentally observed differences in the response to stimuli between clones, 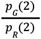 for G vs R and 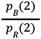 for B vs R, computed for each randomization. The result is shown in **Figure 4H** for TNF-α and in **Figure 5J** for IL-1*β*; examples of trajectories whose ratios in the response match the experimentally observed ones are provided in **Figure S4I**.

#### Predicting the effect of modulating the amount of NF-κB, activating complexes and Nfkbia transcription

We also use our model to test in general the effect of the variation in IκBα, the main negative feedback considered for the system between clones by modulating its transcription rate *k*. Hence we calculate the variation on *k* together with the parameters *C* and *N*_0_, so that the differences in the NF-κB response and in *Nfkbia* expression between clone B and R are reproduced for all the random parameter combinations considered, using again Matlab optimization tools. In particular, we used randomizations to obtain different prototypical parameters of R, *p* _*R*_, as described above, and for each of them generate new parameters sets i.e.: *p* _*B*_ that differ only in the NF-κB amount, the amount of complex and transcription rate,

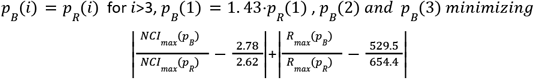

**Figure 6C** shows how these experimentally-constrained change in fitting of *k* reproduces simultaneously the experimental ratio of the expression of *Nfkbia* between clones and responses (derived from **Figure S2E** and **Figure 6B**, respectively). Notice then that for each randomization we generate a new parameter set for clone B differing only in three parameters in an experimentally-constrained way. The predicted resulting changes in the AUC, using the parameters satisfying the equations above, are shown in **Figure 6E**. By applying these three corrections the change in AUC matches better the differences in AUC observed in the experiments, contrarily to simply changing NF-κB amount alone or in combination with the number of activating complexes to fit the response. **Figure 6F** shows instead the change in the number of peaks predicted when the three parameters are changed as prescribed, and for 50% of our “oscillatory simulations” (determined by a number of peaks greater or equal than 2) oscillatory peaks are lost.

#### Mimicking the effect of the ASOs in NF-κB dynamics

To simulate this, we perform different randomization of the parameters of the model for R and then vary the expression of *Nfkbia*, that we assume to be the only quantity affected by ASOs, to match the effect on clone R observed in **Figure S6E** by adjusting properly one parameter: the transcription rate *k*. The predicted effects (fold change in the maximum value and in the AUC) is then evaluated and found to match the experimentally observed one for most of the parameters combination chosen (**Figure 6J**).

### Downmodulation of IκBα using antisense oligonucleotides

The antisense oligos to target the *Nfkbia* gene were designed by Aum Biotech, LLC (Philadelphia, PA, USA). Four custom antisense oligos (in the final concentration of 5 μM) and scrambled control (5 μM) were used to treat MEFs for 24 hours to reduce the expression of IκBa.

## QUANTIFICATION AND STATISTICAL ANALYSIS

We utilized Matlab, R and GraphPad Prism for analysis of the data. We used Student’s t test for comparing two groups and multiple comparisons were done through Anova, Kruskal-Wallis and Mann-Whitney tests as described in each corresponding figure legend. Boxplots represent median, 25th and 75th percentiles, while whiskers cover 2.7 times the standard deviation. Error bars are presented as mean ± SD. p values *p<10^−2^, ** p<10^−3^, *** p<10^−4^ were considered as statistically significant.

## SUPPLEMENTARY MATERIAL

## Tables

## Supplementary movies

Movie S1: Dynamics of the pool of MEFs upon 10 ng/ml TNF-α.

Movie S2: Dynamics of clone B upon 10 ng/ml TNF-a

Movie S3: Dynamics of clone R upon 10 ng/ml TNF-a

Movie S4: Dynamics of clone G upon 10 ng/ml TNF-a

Movie S5: Dynamics of clones B upon 100 ng/ml IL-1*β*

Movie S6. Dynamics of clones G upon 100 ng/ml IL-1*β*.

Movie S7: Dynamics of clone B treated with either mock or *Nfkbia* targeting ASOs upon 10 ng/ml TNF-α.

Movie S8. Dynamics of clone R treated with either mock or *Nfkbia* targeting ASOs upon 10 ng/ml TNF-α.

## Supplementary figure captions

**Figure S1.**
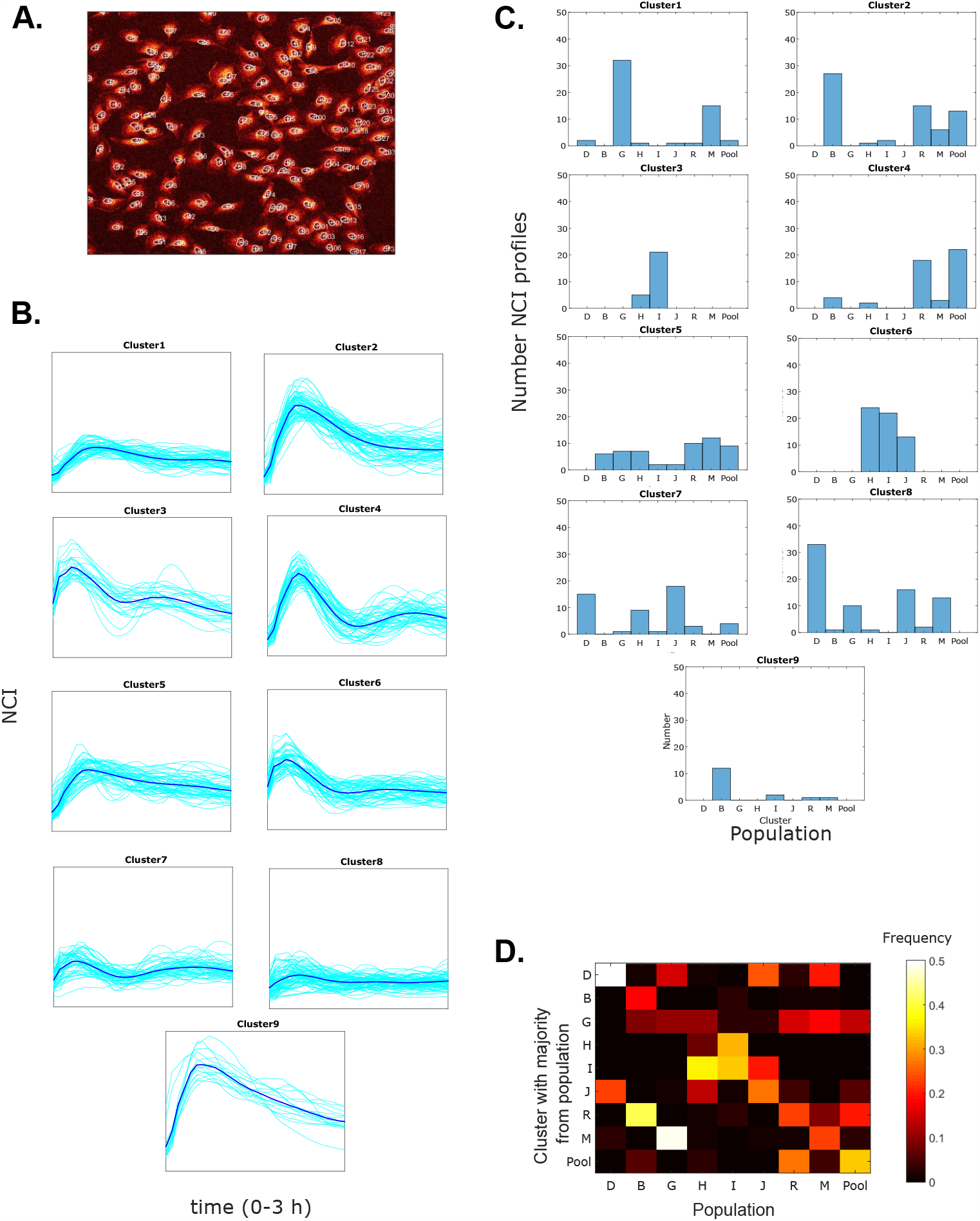
**A**. Example of nuclei detected using our custom-made software for cell segmentation and tracking in a typical image of the 750μm x 750μm field of view. **B**. A realization of our k-means clustering strategy (k=9), where NF-κB dynamic profiles of cells from the 8 clonal populations and the pool are clustered according to their shape in 9 clusters. **C**. Example of the histogram with the number of cells of each population observed in each cluster generated in S1B. **D**. Example of the computed probability of co-clustering of NF-κB dynamic profiles of each population in a given cluster in a single realization of the stochastic clustering algorithm.

**Figure S2.**
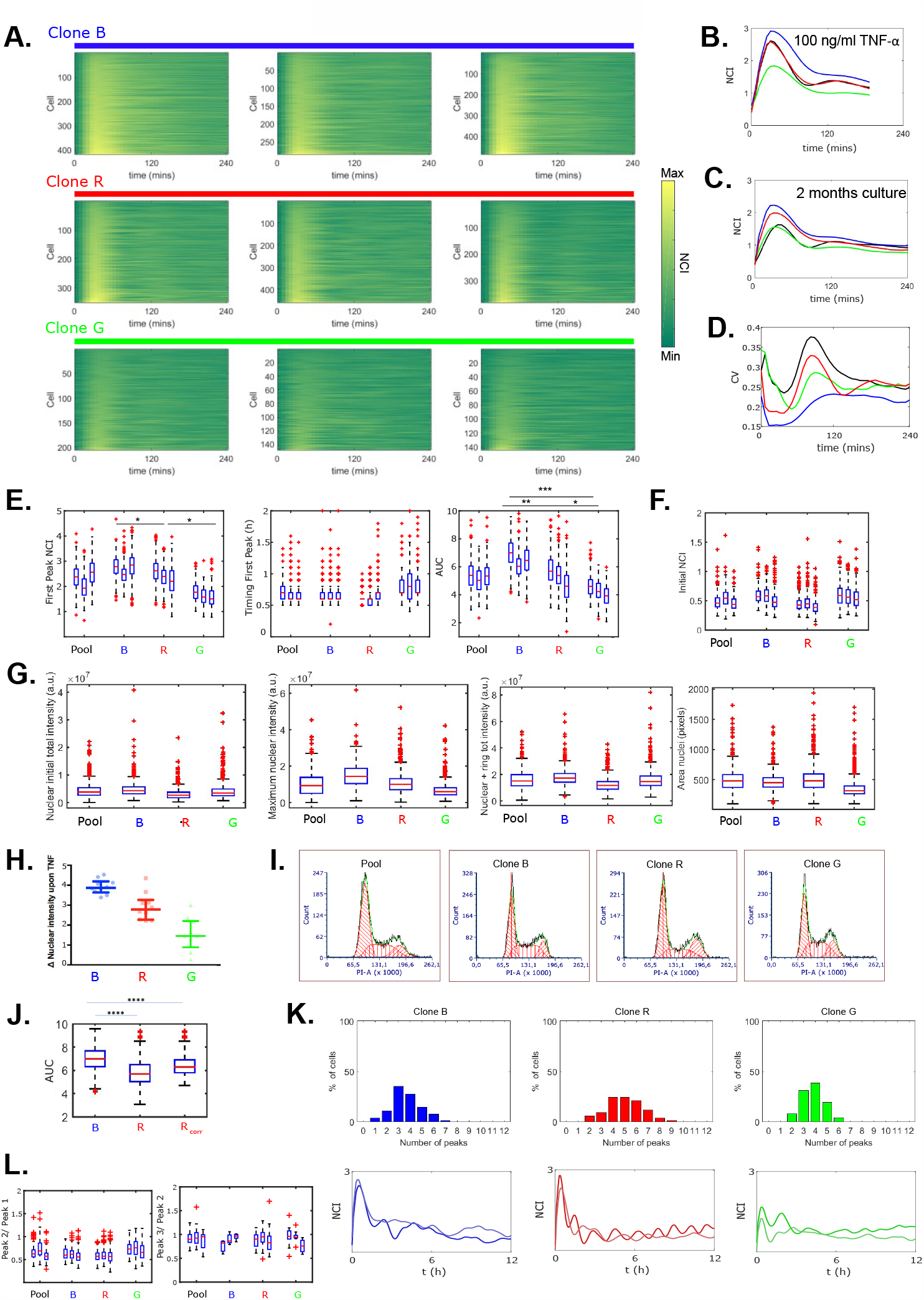
**A**. Dynamic heatmap of the response of the clones B, R and G to 10 ng/ml TNF-α in biological triplicates. **B**. Average NF-κB dynamic profiles of pool and the three clones upon 100 ng/ml TNF-α. **C**. Average NF-κB dynamic profiles of the pool and the three clones B, R, G kept in culture for 8 weeks. **D**. Coefficient of variation (standard deviation divided by the mean) of the pool and the three clones. **E**. Dynamical features of the response to TNF-α in three biological replicates: value of the first peak, timing and area under the curve (AUC). **F**. Average basal initial NCI value (before stimulation) for the three clones. **G**. Features of NF-κB dynamics of the pool and three clones quantified by absolute NF-κB intensities. **H**. Maximum response to TNF-α computed by manually segmenting the nuclei. **I**. Estimation of the cell cycle phases of each population by FACS analysis. **J**. Values of the AUC obtained for clone B, clone R, and clone R by correcting in silico for the differences in cell cycle between B and R. **K**. Distribution of the number of peaks in the three clones for 12 hours of TNF-α stimulation (Top), 12 hour-long time series representative of the heterogeneous oscillatory dynamics observed within the clonal populations (Bottom). **L**. Ratios of the peak values for different peak numbers calculated in three experimental replicates per population, *p<10^−2^, ** p<10^−3^, *** p<10^−4^, multiple comparisons through Kruskal-Wallis.

**Figure S3.**
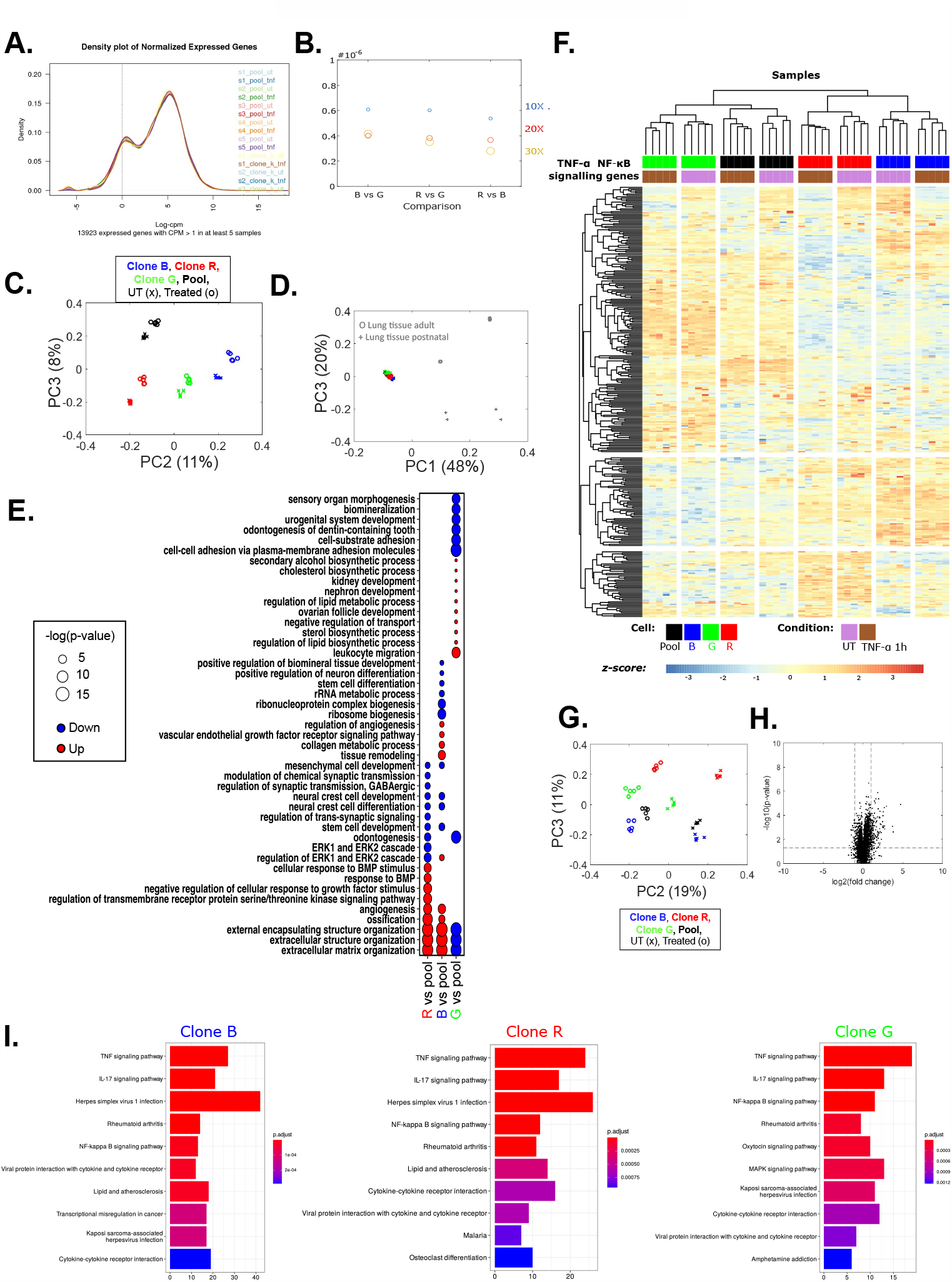
**A.** Density plot of normalized expressed genes for each sample. **B**. Estimated frequency of substitution between clones calculated by the ratio of the number of SNPs and the length of the mapped genome. Increasing marker sizes indicate more stringent criteria for SNPs calling (5x, 10x and 20x coverage). **C**. Additional dimensions of the PCA for our samples’ transcriptomes (% of the variance indicated in parenthesis). **D**. PCA of samples from our clones performed jointly with transcriptomic data from two other mouse tissues (% of the variance indicated in parenthesis). **E**. Gene ontology analysis of the differentially expressed genes between the UT samples of each clonal population and the pool. **F**. Hierarchical clustering of the genes from the “TNF-α NF-κB signaling pathway (Mus musculus)” list from *Wikipathways*. **G**. Additional dimensions of the PCA when considering only NF-κB targets (% of the variance indicated in parenthesis). **H**. Volcano plot for the transcriptomic data of the pool. **I**. KEGG enriched pathways when looking at genes up/down regulated upon TNF-α in the indicated clones.

**Figure S4.**
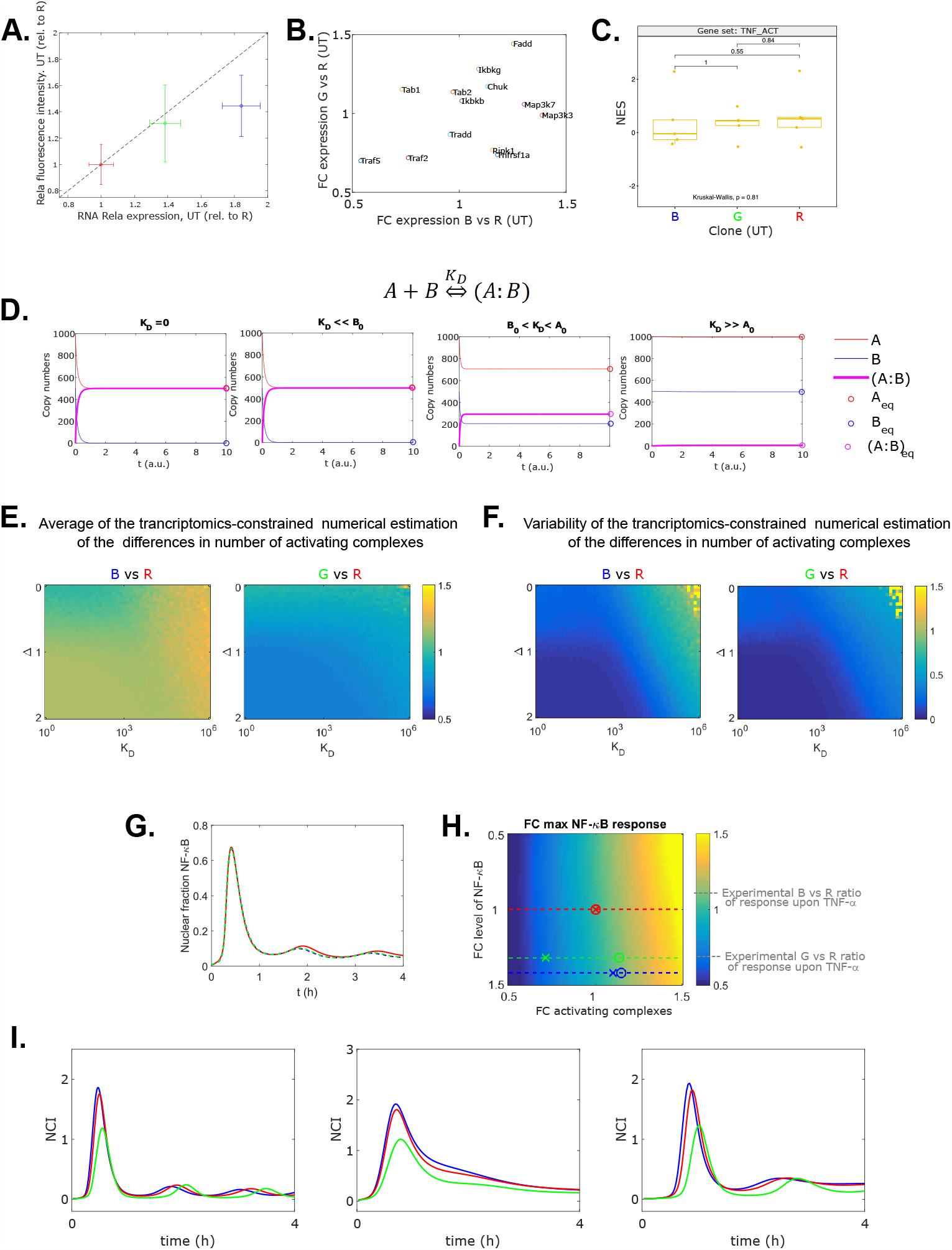
**A**. Expression of RelA gene assessed by RNA-seq plotted against the average fluorescent intensity of the protein per cell in each clonal population with respect to clone R. Error bars denote standard deviation of each measurement, dashed line represents perfect identity. **B**. Fold change expression of genes in the “TNF-α to NF-κB” list, B vs R against G vs R values. **C**. ssGSEA analysis of the “TNF-α to NF-κB” samples (multiple comparisons, Kruskal-Wallis). **D**. Examples of simulations of association-dissociation reaction dynamics for different values of the dissociation constant (*K*_*D*_); circles indicate equilibria inferred from the equilibrium equations. **E**. Transcriptomic-constrained prediction of the FC in the number of complexes formed upon TNF-α for clone B vs R, and for G vs R, for different values of Δ and *K*_*D*_. **F**. Variability in our transcriptomic-constrained prediction of the simulations of E., computed as the standard deviation of the simulations. **G**. Numerical simulation with our mathematical model of NF-κB dynamics of the clone R (red line) and varying only the NF-κB amount to mimic the experimentally observed higher values of clone B and G (dashed blue and green, respectively). **H**. Numerical simulation of the variation in the NF-κB response with NF-κB amount and with the number of activating complexes using our mathematical model. Dashed color lines indicate experimentally observed levels of NF-κB for clone B and G, taken R as reference. Crosses indicate combinations reproducing the experimentally observed variation B vs R and G vs R for TNF-a, displayed on the colorbar with gray dashed lines. Circles indicate the same, but for IL-1*β*. **I**. Examples of randomizations where we have reproduced the procedure schematically represented in H, adjusting the number of activating complexes and of NF-κB so the fold change in the responses across clones (first peak NCI) match those observed experimentally (on average).

**Figure S5.**
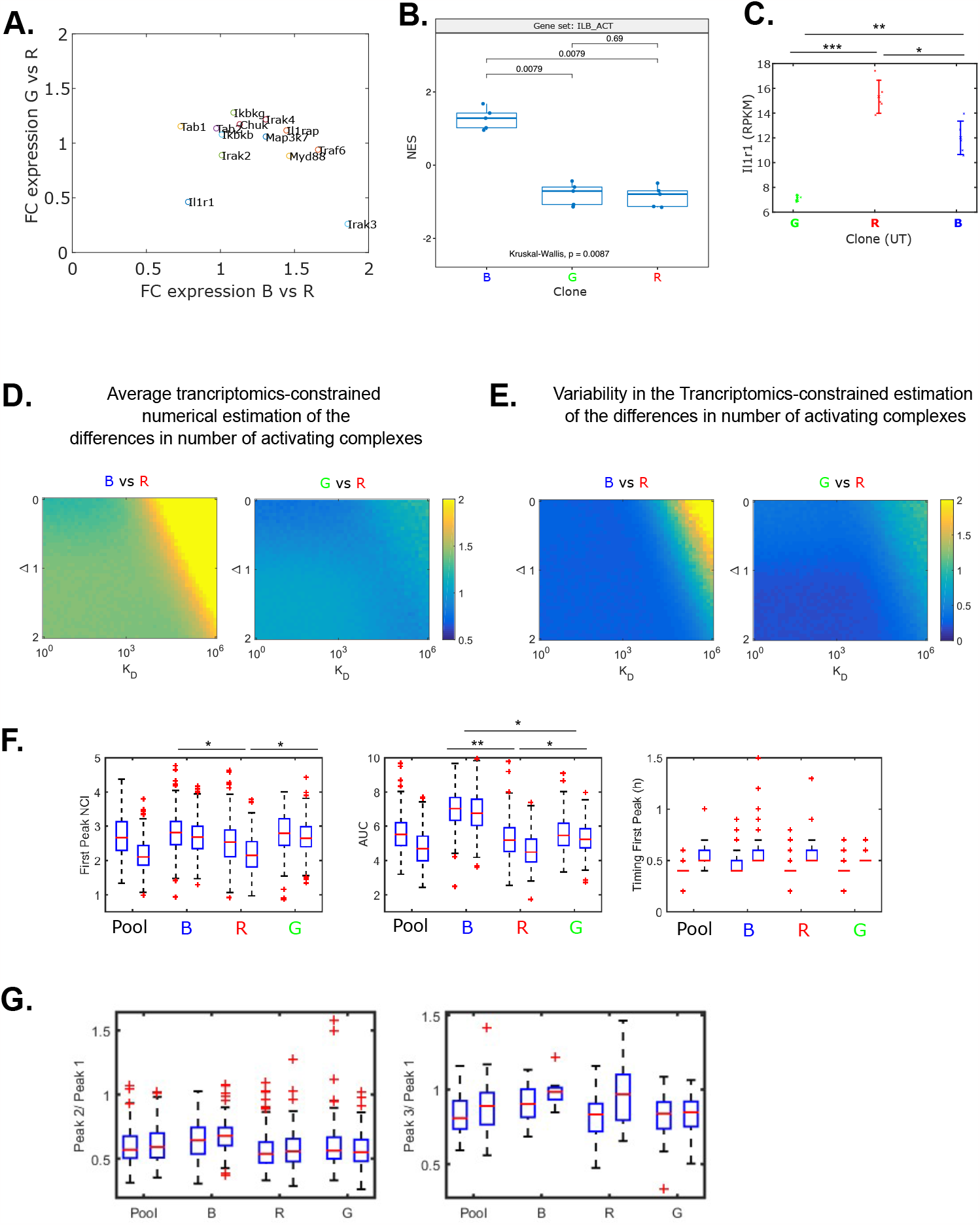
**A**. Fold change expression of genes in the “IL-1*β* to NF-κB” list, B vs R against G vs R values. **B**. ssGSEA analysis of the “IL-1*β* to NF-κB”, only samples from B show statistical difference with the rest. **C**. Expression of *Il1r1* across the clones. **D**. Transcriptomic-constrained prediction of the number of complexes formed upon IL-1*β* for clone B vs R, and for G vs R, for different values of Δ and maximum values of *K*_*D*_ considered. **E**. Variability in our transcriptomic-constrained prediction shown in C., computed as the standard deviations of the simulations performed for each pair of *K*_*D*_ considered and Δ. **F**. Dynamical features of the response to IL-1*β* in two biological replicates. **G**. Features of the oscillatory peaks across the populations of clones upon IL-1*β*. *p<10^−2^, ** p<10^−3^, *** p<10^−4^, multiple comparisons through one way Anova (panel C) and Kruskal-Wallis (remaining panels).

**Figure S6.**
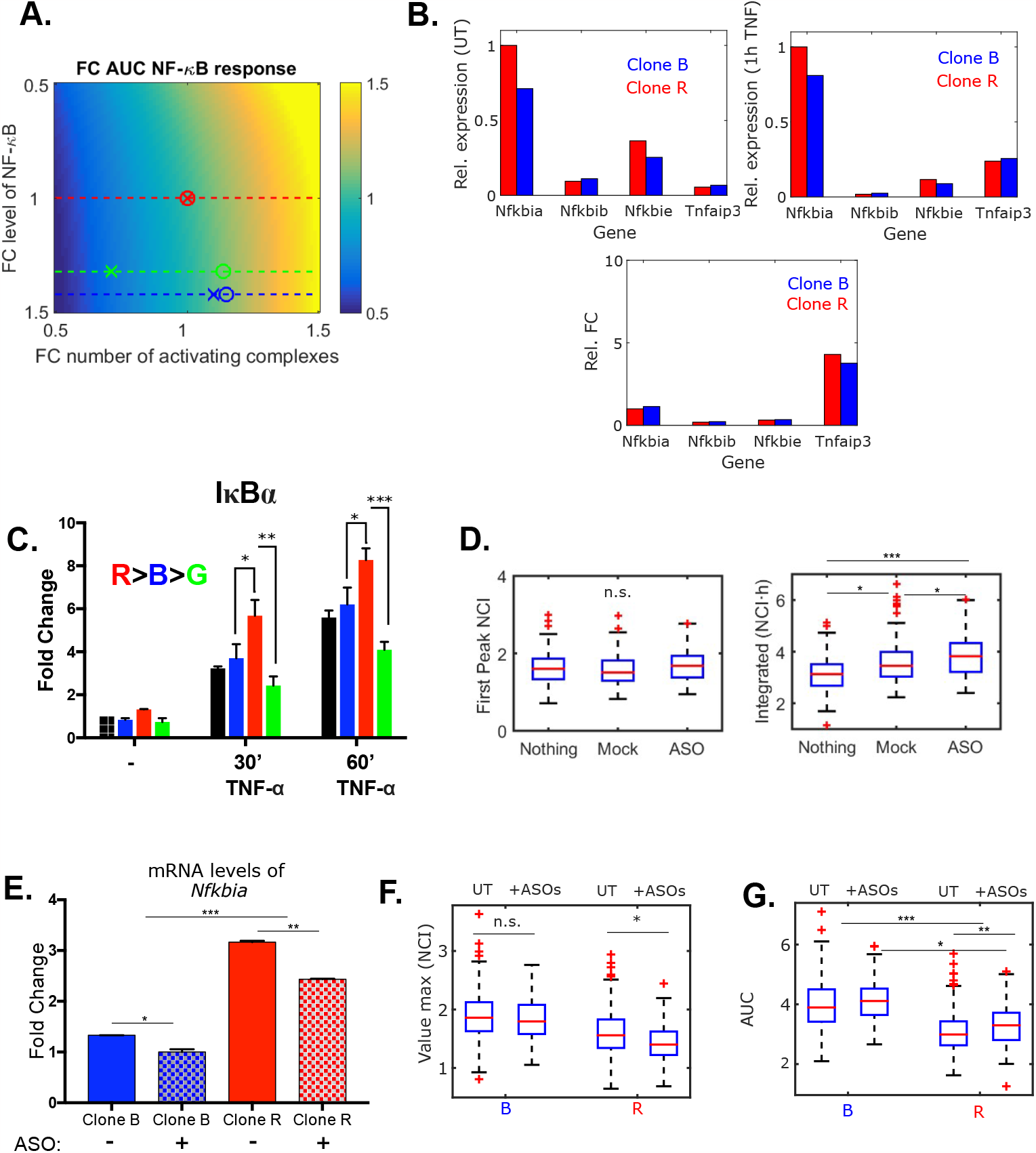
**A**. Numerical simulation for the parameter set of Table 3 of the variation in the AUC of the NF-κB dynamics with the NF-κB amount and the number of activating complexes. Dashed lines indicate experimentally observed levels of NF-κB for clone B and G, taken R as reference. Crosses indicate combinations reproducing the change in the early NF-κB responses respect to R observed for TNF-a, circles for IL-1*β*. **B**. From left to right: expression for UT samples, samples treated with TNF-α for 1h, and FC expression of the negative feedbacks with respect to the values of *Nfkbia* for clone R. **C**. Expression levels (fold change respect UT pool) of IκBα by RT-qPCR. Error bars show the standard deviation of replicates. **D**. Effect of the scrambled (MOCK) ASOs and *Nfkbia-*targeting ASOs on the maximum NCI and AUC of the pool population in comparison to the untreated condition (Nothing). **E**. Relative mRNA levels of IκBα after 24 hours of *Nfkbia-*targeting ASO treatment. Error bars show the standard deviation of replicates. **F**. Quantification of the maximum response and **G**. the AUC for untreated and *Nfkbia-*targeting ASO treated cells upon TNF-α in a biological replicate of the experiment shown in Figure 6. *p<10^−2^, ** p<10^−3^, *** p<10^−4^, multiple comparisons through Kruskal-Wallis except in panel B and E, where two-way Anova was used.

